# Human patient-specific FOXG1 syndrome mouse model revealed FOXG1-MYCN-mediated regulation of protein homeostasis in neurodevelopmental disorder

**DOI:** 10.1101/2025.09.27.678882

**Authors:** Shin Jeon, Liwen Li, Ji-Hwan Moon, Dongjun Shin, Jaein Park, Eunjin Kwak, Jae W. Lee, Soo-Kyung Lee

## Abstract

Neurodevelopmental disorders are characterized by disruptions in brain development, resulting in cognitive, behavioral, and neurological impairments. FOXG1 syndrome (FS), caused by heterozygous mutations in the *FOXG1* gene, exemplifies a severe monogenic neurodevelopmental disorder. To investigate its pathogenesis, we generated a patient-specific W300X mouse model carrying a truncation variant of *FOXG1*. We found that the truncated FOXG1 protein in W300X-heterozygous (W300X-Het) mice is more abundant and more nuclear-localized than the full-length FOXG1 protein, implicating a pathogenic mechanism involving the truncated protein. Interestingly, W300X-Het mice exhibited profound abnormalities in the dentate gyrus, including disrupted neurogenesis, impaired granule cell migration, and altered dendritic morphology. Transcriptomic profiling identified broad dysregulation in protein homeostasis pathways, particularly ribosomal biogenesis, translation, and proteostasis. Disruption of the FOXG1-MYCN pathway, critical for robust protein synthesis during neural stem cell division, synaptogenesis, and synaptic plasticity, emerged as a key mechanism underlying these defects. In parallel, microglial activation and inflammation were markedly increased in the dentate gyrus, contributing to a pro-inflammatory environment that exacerbates neurogenic and structural deficits. Consistent with hippocampal dysfunction in FS patients, W300X-Het mice exhibited significant spatial learning and memory impairments. Together, our study highlights disrupted protein homeostasis and neuroinflammation as key drivers of FS pathogenesis, providing a framework for developing therapeutic strategies targeting these pathways.

## INTRODUCTION

Neurodevelopmental disorders (NDDs) represent a diverse group of conditions marked by disrupted brain development and function, manifesting as cognitive, behavioral, and neurological impairments. Advances in genetic diagnostics have enabled the identification of numerous monogenic NDDs, where a single gene variant underlies complex neurological and behavioral deficits. Studying the mechanisms of these monogenic disorders offers critical insights into shared clinical symptoms and the genetic networks that govern brain maturation and function.

Heterozygous mutations in the *FOXG1* gene cause FOXG1 syndrome (FS), a severe neurodevelopmental disorder characterized by intellectual disability, epilepsy, and motor impairments^1-5^. As a monogenic disorder, FS provides a unique opportunity to uncover pathogenic mechanisms underlying a range of NDDs, given the pivotal role of FOXG1 in orchestrating the gene regulatory networks essential for forebrain development and maturation. During neural tube patterning, FOXG1 expression is specifically induced in the forebrain, where it governs neural stem cell (NSC) proliferation and differentiation, neuronal migration, axon projection, and dendrite morphogenesis^6-11^. FOXG1 executes these diverse functions as a transcription factor (TF) by regulating target genes in a cell context-specific manner^10,12^.

Among the gene variants causing FS, truncation mutations, such as nonsense and frameshift variants, are the most common^3,4,13,14^. Variants occurring downstream of the DNA-binding domain (DBD) are of particular interest, as they can produce truncated FOXG1 proteins retaining the DBD, potentially exerting gain-of-function effects. Addressing this critical aspect of FS pathogenesis necessitates a patient mutation-mimicking mouse model, as existing *Foxg1* heterozygous (*Foxg1^+/-^*) mice are insufficient for studying the specific effects of truncation mutations.

The hippocampus, a brain region integral to learning, memory, and spatial navigation, is frequently implicated in NDDs^15,16^. Structural and functional abnormalities in the hippocampus are hallmark features of conditions such as autism spectrum disorder (ASD), schizophrenia, and epilepsy^17-19^. Within the hippocampus, the dentate gyrus (DG) plays a pivotal role in adult neurogenesis, pattern separation, and the modulation of excitatory input to the hippocampal circuit^20^. Decreased proliferation and differentiation of NSCs in the DG have been observed in animal models of many NDDs, such as ASD, Rett syndrome, Down syndrome, and Fragile X syndrome, correlating with deficits in memory and social behaviors^17,21^. Advances in understanding DG-specific pathophysiology will provide crucial insights into the mechanisms underlying hippocampal dysfunction in NDDs and offer potential therapeutic targets.

FOXG1 plays a pivotal role in hippocampal development and maturation. FOXG1 regulates hippocampal specification by modulating the boundaries of the cortical hem, a key signaling center adjacent to the hippocampus primordium, during embryonic development^22^. Conditional deletion of *Foxg1* in the postnatal dentate gyrus (DG) results in impaired granule cell maturation and reduced mossy fiber connectivity^23^. Conversely, overexpression of FOXG1 in rapidly dividing neural progenitor cells (NPCs) in the adult DG leads to reduced survival, impaired maturation, and abnormal morphology of newborn neurons^24,25^, underscoring the importance of precise FOXG1 dosage for DG neurogenesis. *Foxg1^+/−^* mice exhibit deficits in the generation and survival of DG neurons, with impairments becoming more pronounced with age, while embryonic DG neurogenesis remains relatively intact^26^. Interestingly, despite significant postnatal cell loss in the DG, apoptosis was not increased in *Foxg1^+/−^*mice^26^. This suggests that DG cells in these mice are eliminated through non-apoptotic cell death pathways, such as excitotoxicity and necroptosis, all involving inflammation^27,28^. However, the connection between FOXG1 haploinsufficiency and neuroinflammation remains unexplored.

In this study, we generated an FS patient-derived mouse model carrying the W300X truncation mutation to investigate the pathophysiology of FS. The W300X-heterozygous (W300X-Het) mouse model not only accurately mimicked FS genetics but also recapitulated altered hippocampal architecture, a key feature of FS. Our findings revealed that the W300X mutation leads to the production of a truncated FOXG1 protein, which shows molecular characteristics distinct from the full-length FOXG1 protein. W300X-Het mice displayed profound disruptions in the cellular, structural, and functional integrity of the DG, including impaired neurogenesis, aberrant neuronal migration, and altered granule cell organization. Transcriptomic analyses further highlight the widespread dysregulation of synaptic, translational, and proteostatic pathways in the W300X-Het DG, indicating a heightened level of cellular stress. We identified the FOXG1-MYCN pathway as a critical regulatory axis, where FOXG1 collaborates with MYCN to drive the expression of genes essential for ribosomal biogenesis and translation, processes vital for NSC proliferation, synaptogenesis, and synaptic plasticity. Accordingly, W300X-Het mice exhibited microglial activation, inflammation in the DG, and significant learning and memory deficits. Collectively, our study offers new insights into the pathogenesis of NDDs, highlighting disrupted protein homeostasis and the FOXG1-MYCN pathway as key contributors to FS.

## RESULTS

### The generation of nonsense-type FS patient-specific mouse model

As *FOXG1* is a single exon gene, FS-causing nonsense or frameshift variants in *FOXG1* are likely to produce protein products that lack the C-terminal portion of the FOXG1 protein. Notably, 46% of FS cases have gene variants that can produce a truncated form of FOXG1 protein, including 15% nonsense variants^3^. Particularly, 15% of FS variants have mutations after the DBD of FOXG1, which can generate an N-terminal fragment form of FOXG1 that contains the DBD.

To test if the truncated FOXG1 proteins are involved in the pathophysiology of FS, we modeled p.Trp308Ter (p.W308X) resulting from two different nucleotide variants c.923G>A and c.924G>A for animal modeling (Fig. 1a). We generated a knock-in mouse line, in which the W300X mutation (c.900G>A, p.W300X corresponding to the human p.W308X variant) was inserted to the mouse *Foxg1* gene (Fig. 1a,b). W300X homozygous mice (Foxg1^W300X/W300X^, W300X-Hom mice) showed a markedly reduced forebrain and perinatal death with 100% penetrance (Fig. 1c, Supplementary Fig. 1). These results indicate that W300X protein cannot functionally replace full-length FOXG1 protein (referred to as FOXG1-fl) despite the presence of DBD. W300X heterozygous mice (Foxg1^W300X/+^, W300X-Het mice), which accurately recapitulate FS-causing haploinsufficiency genetic conditions, survived into adulthood, allowing us to study the role of the W300X truncated protein in the postnatal maturation of the brain.

**Figure 1.**
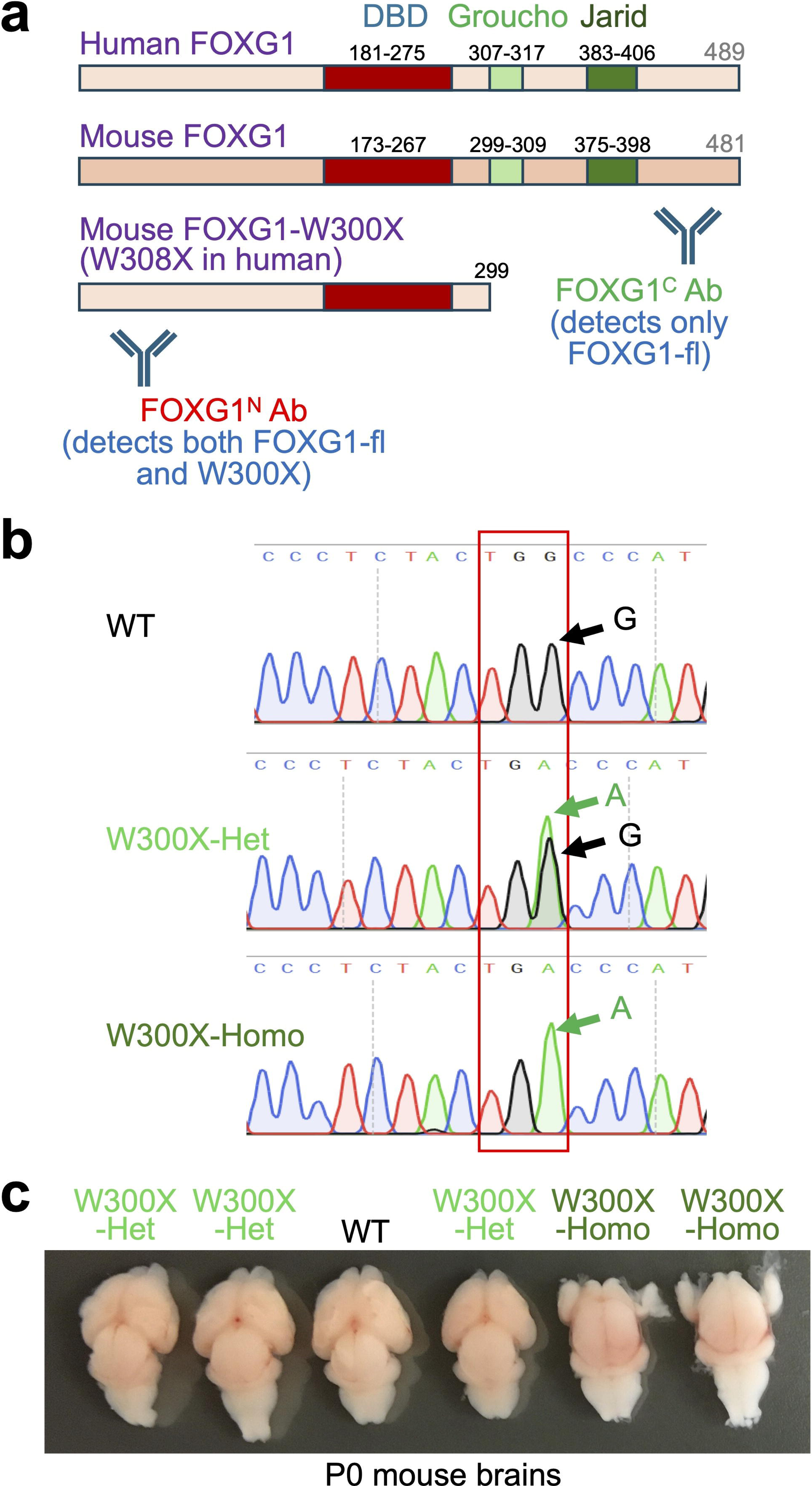
The generation of the W300X mouse model, a nonsense type of FOXG1 patient-specific model. (a) Schematic representation of human FOXG1, mouse FOXG1, and mouse W300X proteins. FOXG1^N^ and FOXG1^C^ antibodies specifically target the N-terminal and C-terminal regions of FOXG1, respectively. (b) Sanger sequencing confirms the presence of the *Foxg1* gene variant c.900G>A (p.W300X) in W300X-Het and W300X-Homo mice, validating the successful establishment of the W300X mouse model. (c) Representative brain images at P0. W300X-Homo mice exhibited severe forebrain hypoplasia compared to wild-type (WT), indicating the loss-of-function nature of the allele.

### Expression of W300X truncated protein exceeds full-length FOXG1 in W300X-Het brains

To evaluate whether the *W300X* mutant allele produces the truncated FOXG1 protein in the brain, we utilized antibodies targeting the N-terminal (FOXG1^N^ Ab) and C-terminal (FOXG1^C^ Ab) regions of FOXG1. FOXG1^N^ Ab recognizes both FOXG1-fl and W300X proteins, whereas FOXG1^C^ Ab detects only FOXG1-fl due to the absence of the C-terminal region in the W300X protein (Fig. 1a).

We dissected the cortex and hippocampus from P1 and P21 mice and performed western blot analysis. The FOXG1^C^ Ab detected a prominent ∼70 kDa band corresponding to FOXG1-fl, whose intensity was substantially reduced in W300X-Het brains relative to WT (Fig. 2a), establishing that the functional FOXG1-fl protein was decreased in W300X-Het brains. Interestingly, additional smaller molecular weight bands were observed with FOXG1^C^ Ab, indicating that multiple forms of FOXG1’s C-terminal fragments are expressed in the brain (Fig. 2a).

**Figure 2.**
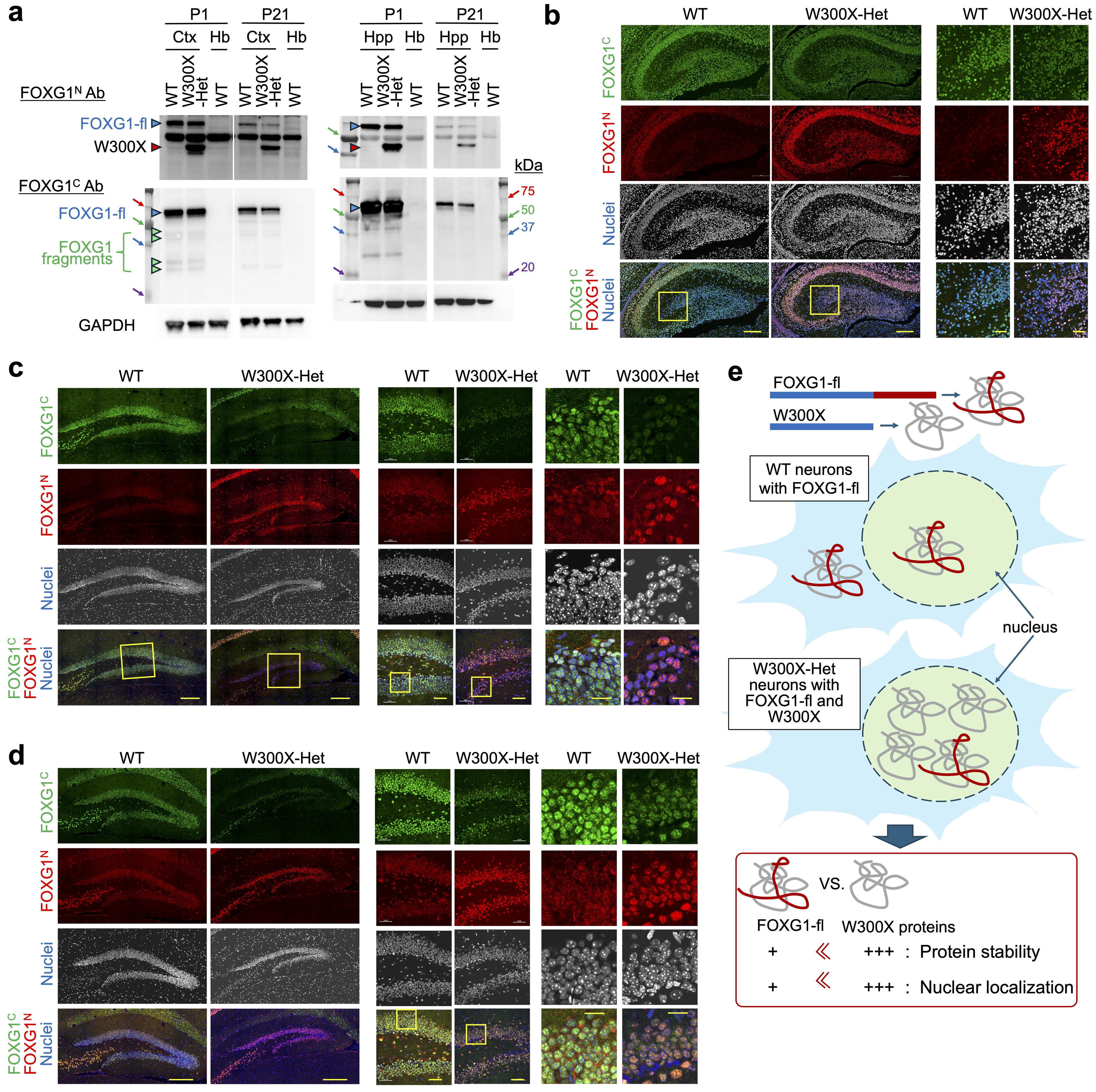
W300X truncation leads to altered FOXG1 protein abundance, stability, and localization in hippocampus. (a) Western blot analysis of FOXG1 protein levels in the cortex (Ctx), hippocampus (Hpp), and hindbrain (Hb) at P1 and P21. W300X denotes samples from W300X-Het brains. The hindbrain served as a negative control due to the absence of FOXG1 expression. Full-length FOXG1 (FOXG1-fl) levels were significantly reduced in the W300X-Het cortex and hippocampus compared to corresponding WT tissues. Notably, the W300X fragment was detected exclusively in W300X-Het samples by the FOXG1^N^ antibody but not by the FOXG1^C^ antibody, indicating the lack of the C-terminal region. (b-d) Immunostaining analysis of hippocampal regions at P1 (b), P21 (c), and adulthood (d) using FOXG1^C^ (green) and FOXG1^N^ (red) antibodies. In the W300X-Het hippocampus, FOXG1-fl detected by the FOXG1^C^ antibody was reduced, whereas the combined signals of FOXG1-fl and W300X proteins, recognized by the FOXG1^N^ antibody, were enhanced. Additionally, the FOXG1^N^ antibody revealed predominantly nuclear signals in W300X-Het brains, whereas it exhibited a more diffuse distribution in WT brains. Each panel consists of paired columns (WT and W300X-Het) showing images at the same magnification. Magnified regions are highlighted by yellow rectangles. Scale bars: 200 μm (left panels in b and d), 100 μm (left panel in c), 50 μm (right panel in b, middle panels in c and d), 20 μm (right panels in c and d). (e) Model of molecular behavior of FOXG1-fl and W300X proteins. The W300X truncation lacks the C-terminal region (red portion of FOXG1-fl). In W300X-Het neurons, the total level of FOXG1-fl protein is reduced compared to WT, while W300X protein levels exceed those of FOXG1-fl. In WT neurons, FOXG1-fl is distributed in both nucleus and cytoplasm. In W300X-Het neurons, both FOXG1-fl and W300X proteins show nuclear-predominant localization, with the W300X protein exhibiting greater stability and nuclear enrichment relative to FOXG1-fl.

Notably, the FOXG1^N^ Ab identified two distinct bands in W300X-Het samples: ∼70 kDa FOXG1-fl band and ∼40 kDa band corresponding to the truncated W300X protein (Fig. 2a). In contrast, in WT samples, only the FOXG1-fl band was present. These results indicate that the *W300X* allele produces the truncated FOXG1 protein W300X, which lacks the C-terminal region but retains the N-terminal domain (Fig. 1a). Strikingly, in W300X-Het brains, the truncated W300X band appeared more intense than the FOXG1-fl band, despite the 1:1 allele ratio, indicating that the mutant protein accumulates to higher levels than FOXG1-fl (Fig. 2a). These results suggest that W300X protein is more stable than FOXG1-fl (schematics in Fig 2e).

To further assess the relative levels of FOXG1-fl and W300X proteins, we performed immunohistochemistry (IHC) analyses on the hippocampus from P1 to adult stages using FOXG1^N^ Ab and FOXG1^C^ Ab. From the perinatal to adult stages, FOXG1 is expressed in the CA1, CA2, and CA3 pyramidal neurons and dentate gyrus (DG) (Fig. 2b-d, Supplementary Fig. 2). FOXG1^C^ Ab staining revealed a weaker signal in the W300X-Het hippocampus compared to WT controls (Fig. 2b-d), underlining reduced FOXG1-fl protein levels in the mutant mice. Conversely, FOXG1^N^ Ab staining intensity, which reflects the combined level of FOXG1-fl and W300X proteins, was elevated in the W300X-Het hippocampus relative to WT (Fig. 2b-d). The increased FOXG1^N^ Ab staining signal was pronounced in the DG at all three stages. These findings are consistent with the western blot results that the W300X protein is expressed at higher levels than FOXG1-fl protein in W300X-Het brains, suggesting increased stability of the truncated protein. Intriguingly, IHC with the FOXG1^N^ Ab revealed a diffuse localization of the FOXG1 protein across both nuclear and cytosolic compartments in the DG of WT mice (Fig. 2b-d). In contrast, FOXG1^N^ Ab signals exhibited a predominantly nuclear localization pattern in the W300X-Het DG (Fig. 2b-d). This result suggests that the W300X truncated protein is primarily localized to the nucleus and enhances the nuclear retention of the full-length FOXG1 (FOXG1-fl) protein (Fig. 2e).

Besides establishing the expression of the W300X protein, these findings provide crucial insights into the molecular consequences of the *W300X* mutation, which may underlie the phenotypic impairments in this mouse model.

### W300X-Het mice show remarkable morphogenic defects in the DG

The DG undergoes a series of intricate morphogenetic processes during development, which are essential for establishing its characteristic crescent-shaped structure and functional integration within the hippocampal circuitry^29,30^. At P1, where the DG is in an early developmental stage, NPCs divide actively, and the subgranular zone (SGZ) neurogenic niche is not yet fully established. Granule cell precursors are migrating from the primary dentate matrix toward the forming granule cell layer, and many granule cells are still immature, showing morphological features of migrating neurons. Initial synaptic connections begin forming between granule cells, mossy fibers, and input from the entorhinal cortex. At P21, the DG reached a more advanced stage of development but still exhibits higher neurogenesis levels than adults. Over the following weeks, neurogenesis rates decrease, reaching adult baseline levels by P60.

DAPI nuclear staining revealed the overall structures of the DG. At P1, relative to WT, the DG primordium in W300X-Het mice was relatively similar in size but exhibited less-defined morphology, particularly at the tip of the upper (suprapyramidal) blade, which appeared blunted and lacked the sharp contour (Fig. 2b).

By P21, the W300X-Het DG showed pronounced structural abnormalities. The total length and thickness of the upper and lower (infrapyramidal) blades were markedly reduced in W300X-Het mice relative to WT (Fig. 2c). These findings suggest that the developmental trajectory of the DG is severely disrupted in the W300X-Het hippocampus.

In adults, the DG was markedly shorter and thinner in W300X-Het mice than WT (Fig. 2d), suggesting a severe disruption in structural integrity and overall neuronal population maintenance.

### W300X-Het brains exhibit defects in the cellular, structural, and functional integrity of the DG

To assess how the W300X mutation impacts cellular populations and the structural organization of the DG, we performed IHC with a panel of markers delineating various DG cell populations in adult mice. Immunostaining for GFAP and SOX2, markers of NSCs, demonstrated a significant reduction in SOX2^+^GFAP^+^ NSCs in the SGZ of the W300X-Het DG (Fig. 3a). The number of NSCs was markedly reduced in the W300X-Het hippocampus. Further, GFAP^+^ radial glial scaffolds, which typically extend vertically from the SGZ to the molecular layer (ML), were disrupted (Fig. 3a), indicating compromised NSC niche function and impaired radial glia-mediated neuronal projection and migration.

**Figure 3.**
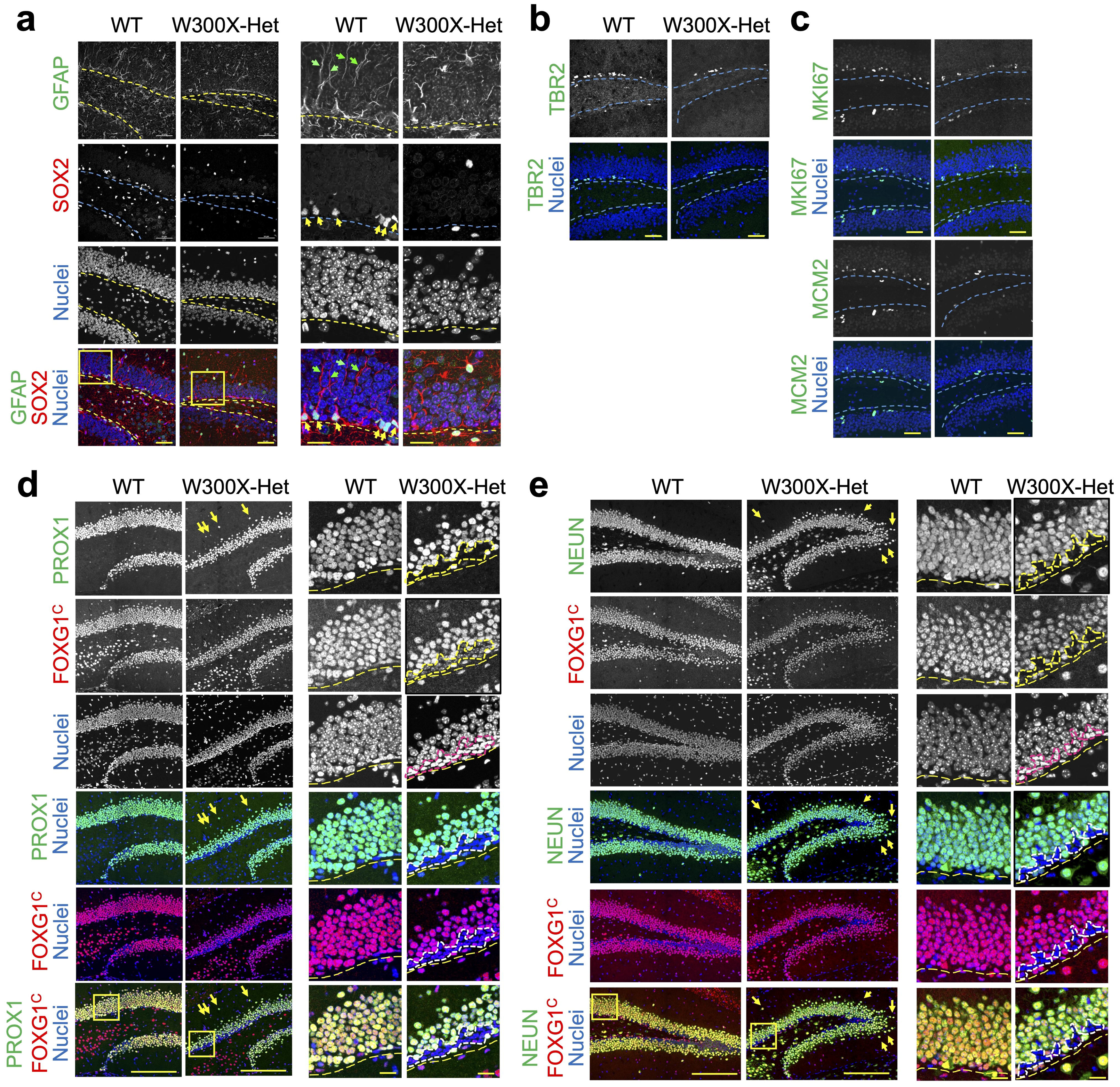
Neural stem cell (NSC) depletion and disorganized granule cells in the dentate gyrus (DG) of W300X-Het brains. (a-e) Immunostaining analysis of the adult DG (a) SOX2^+^GFAP^+^ NSCs (yellow arrows) in the subgranular zone (SGZ) are significantly reduced, and GFAP^+^ radial glial scaffolds (green arrows) are disorganized in W300X-Het mice. (b) TBR2^+^ IPCs are decreased, reflecting impaired NSC lineage progression. (c) MCM2^+^MKI67^+^ proliferative cells are reduced, indicating compromised cell proliferation. (d-e) PROX1^+^NEUN^+^ granule neurons are decreased, disorganized, and displaced outside the compact granule zone (GZ). Yellow arrows denote displaced PROX1^+^NEUN^+^ granule cells mislocalized outside the densely packed granule zone. An aberrant population of PROX1^-^, NEUN^-^, and FOXG1-negative cells with horizontally elongated nuclei (outlined by dotted lines) appears at the SGZ, representing a novel W300X-associated phenotype. Each panel consists of paired columns (WT and W300X-Het) showing images at the same magnification, with magnified regions highlighted by yellow rectangles. Scale bars: 200 μm (left panels in d and e), 50 μm (right panel in a, panels in c and d), 20 μm (right panels in a, d, and e).

The W300X-Het DG exhibited a dramatic reduction in TBR2^+^ intermediate progenitor cells (IPCs) within the SGZ (Fig. 3b), highlighting disrupted neurogenic lineage progression, likely due to reduced NSC proliferation and differentiation. This was further corroborated by significantly fewer MCM2^+^ and MKI67^+^ proliferating cells in the W300X-Het SGZ (Fig. 3c), indicating a severe decline in cellular proliferation.

PROX1^+^ cells, encompassing developmental and mature granule cells, were substantially reduced in the W300X-Het DG (Fig. 3d). In the WT upper blade, PROX1^+^ granule neurons form 8-10 densely packed cell rows within the GZ (granule zone), while in the W300X-Het upper blade, only 4-5 cell rows were observed. Furthermore, PROX1^+^ cells in W300X-Het were less densely packed, with many cells displaced outside the compact GZ (yellow arrows, Fig. 3d). NEUN^+^ mature neurons mirrored this disorganization, with reduced neuronal cell density and mislocalized DG neurons outside the GZ (yellow arrows, Fig. 3e), suggesting defects in cell-cell interactions, neuronal migration, and granule cell organization.

In the adult DG of WT mice, most cells express PROX1 and NEUN, except for a small subset of NSCs and IPCs in the SGZ (Fig. 3d,e). Notably, the W300X-Het SGZ exhibited an aberrant layer of PROX1-, NEUN- and FOXG1-negative cells with horizontally elongated nuclei, distinct from those observed in the WT SGZ (outlined by dotted lines, Fig. 3d,e), indicating a novel phenotype associated with the W300X mutation.

These findings collectively reveal that the W300X mutation disrupts NSC maintenance, impairs neurogenic lineage progression, and causes defects in neuronal migration and granule cell organization in the DG. The emergence of aberrant PROX1/NEUN/FOXG1-negative cells further highlights unique pathological features in the W300X-Het DG.

### W300X-Het brains display deficits in neuronal generation, positioning, and dendrite development

During neurogenesis in the DG, newly generated neurons originate in the SGZ, where their cell bodies remain aligned. To evaluate neurogenesis, we labeled newborn neurons with DCX antibodies^31^. At P40, an adolescent stage characterized by active neurogenesis, DCX^+^ neurons formed a distinct layer along the SGZ (Fig. 4a). This pattern persisted into adulthood, although the rate of neurogenesis declined significantly at this stage (Fig. 4b). In the W300X-Het DG, gaps were evident in the DCX^+^ newborn neuron layer in the SGZ, particularly in the lower blade at P40 (yellow parentheses, Fig. 4a), indicating a stark reduction in neurogenesis. Additionally, many DCX^+^ neurons appeared displaced into the granular zone GZ and molecular layer ML (arrows, Fig. 4a). By adulthood, the reduction in DCX^+^ newborn neurons became more pronounced in the W300X-Het DG, with only sparse DCX^+^ neurons in the lower blade of the W300X-Het DG (yellow parentheses, Fig. 4b). In contrast, the WT DG maintained a continuous layer of DCX^+^ neurons in the SGZ of both the upper and lower blades (Fig. 4b). Furthermore, in the W300X-Het DG, some DCX^+^ neurons delaminated from the SGZ and were displaced into the hilus (magenta arrows, Fig. 4b), further highlighting defects in the granular cell organization.

**Figure 4.**
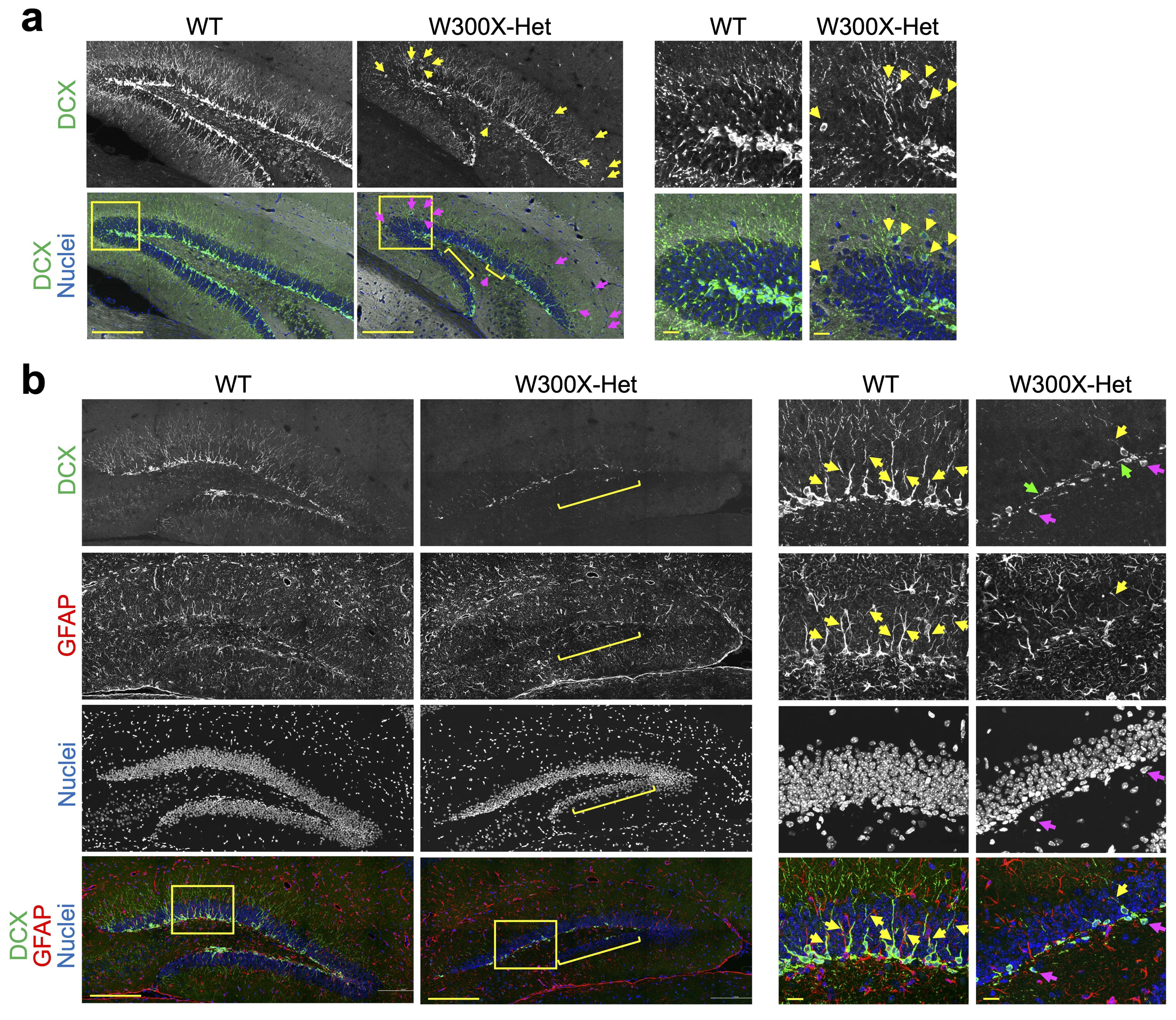
The dentate gyrus (DG) of W300X-Het mice exhibits severe deficits in neurogenesis and dendritic development. (a,b) Immunostaining analysis of the DG at P40 (a) and adulthood (b). (a) Relative to WT, W300X-Het DCX^+^ newborn neurons at P40 are sparse, with gaps in the SGZ and frequent displacement into the GZ and ML. Parentheses indicate regions of the SGZ devoid of newborn neurons in the lower blade of the W300X-Het DG. (b) In adults, relative to WT, W300X-Het DCX^+^ neurons remain reduced and mislocalized. Many DCX^+^ dendrites retain a horizontal orientation, failing to reorient vertically and branch within the ML. These defects correlate with disrupted GFAP^+^ radial glial scaffolding, suggesting impaired guidance for dendritic growth and synaptic integration. Yellow arrows mark the close association between DCX^+^ dendritic processes and GFAP^+^ radial glial scaffolds. Green arrows denote DCX^+^ dendrites that retain an abnormal horizontal projection pattern. Magenta arrows indicate DCX^+^ neurons delaminated from the SGZ. Each panel consists of paired columns (WT and W300X-Het) displaying images at the same magnification, with magnified regions outlined by yellow rectangles. Scale bars: 200 μm (left panels in a and b), 20 μm (right panels in a and b).

During their maturation, newborn DCX^+^ neurons in the DG undergo dynamic changes in dendritic morphology^31^. Initially, dendrites grow horizontally within the plane of the SGZ, but as neurons mature, their dendrites reorient and extend vertically toward the ML. Within the ML, dendrites undergo extensive branching, establishing a highly intricate arborization critical for synaptic connectivity (yellow arrows, Fig. 4b). In the W300X-Het DG, dendritic morphology was profoundly disrupted (yellow arrows, Fig. 4b). Many DCX^+^ dendrites retained a horizontal projection pattern, failing to reorient vertically toward the ML (green arrows, Fig. 4b). Even when some dendrites reached the ML, they failed to branch extensively, compromising their ability to form appropriate connections (yellow arrows in W300X-Het, Fig. 4b).

Radial glial cells (RGCs) and molecular signals, such as Reelin and Notch, are essential for guiding dendritic growth, orientation, and complexity, thereby ensuring the integration of newborn neurons into hippocampal circuits^32-34^. GFAP^+^ RGCs in the SGZ extended their processes into the ML in WT, providing structural guides for dendritic orientation and growth (yellow arrows, Fig. 4b). DCX^+^ dendritic processes were closely associated with these GFAP^+^ radial glial scaffolds in the WT DG, demonstrating a robust interaction (yellow arrows, Fig. 4b). However, in the W300X-Het DG, this extensive interaction between dendritic processes and radial glial scaffolds was markedly disrupted (yellow arrows, Fig. 4b). This disruption of radial scaffolding, combined with impaired dendritic development, likely exacerbated the defects in neuronal circuit formation.

Overall, our results demonstrate that dendritic morphogenesis and integration into hippocampal circuits are severely compromised in the W300X-Het brain.

### Transcriptome analysis of W300X-Het hippocampus revealed dysregulation of the genes implicated in neurogenesis, neuronal connection, and synapses

To understand the mechanistic basis underlying the striking cellular defects in the hippocampus of W300X-Het mice, we conducted RNA-seq analysis at the P1 stage. As the W300X-Het hippocampus at P1 has not shown the drastic cellular architecture disruption observed at later stages, P1 represents a window to capture transcriptomic alterations that reflect the molecular perturbations caused by the W300X mutation, before the compounding effects of structural reorganization obscure these early molecular changes. RNA-seq identified 710 differentially expressed genes (DEGs) in the W300X-Het hippocampus relative to WT mice (413 upregulated and 297 downregulated genes, referred to as up-DEGs and down-DEGs, respectively; Fig. 5a, Supplementary Table 1). To delineate the disrupted molecular pathways in the W300X-Het hippocampus, we performed a comprehensive analysis of DEGs using multiple ontological and pathway enrichment databases^35^.

**Figure 5.**
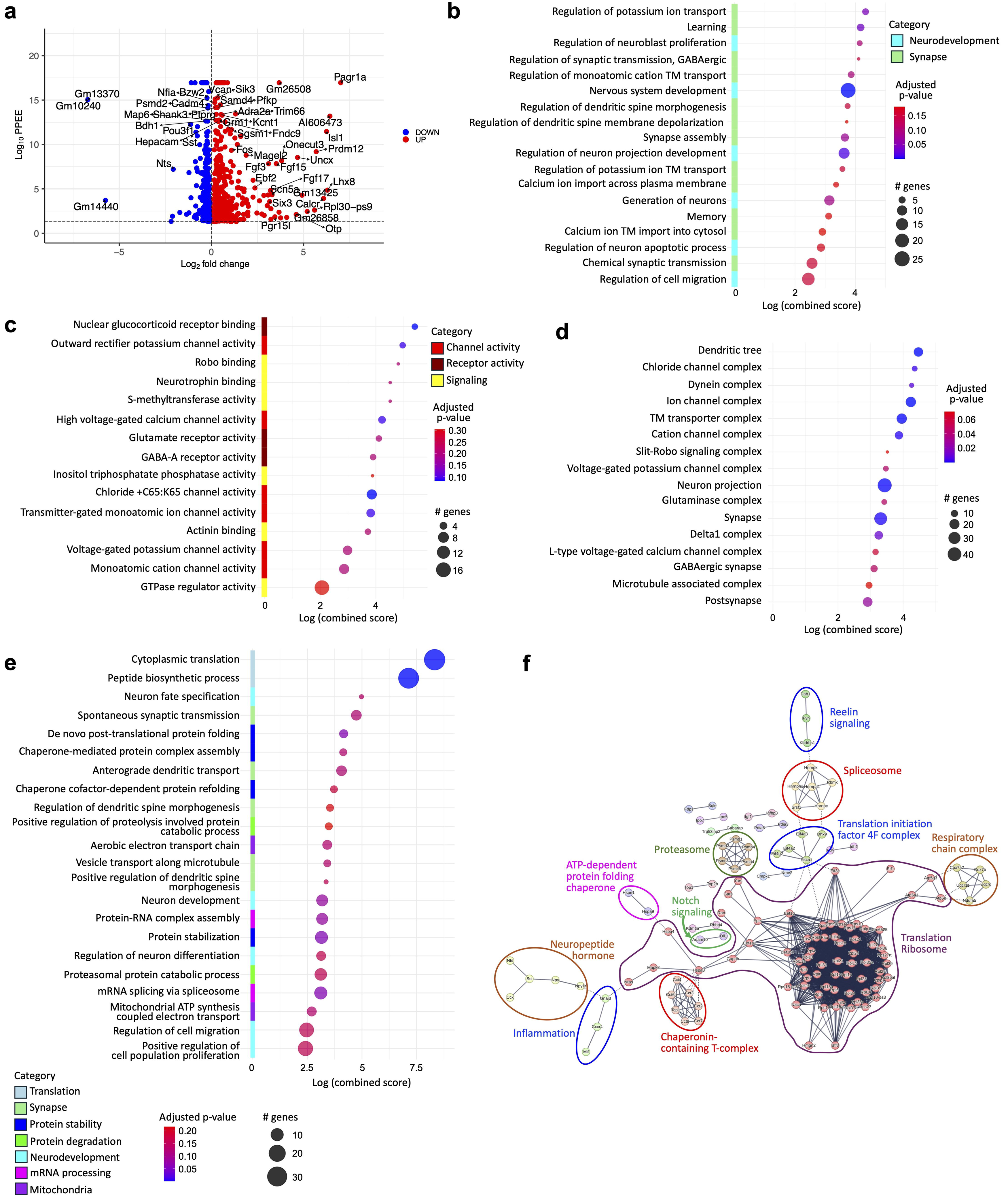
Transcriptomic dysregulation in W300X-Het hippocampus highlights altered synaptic, translational, and proteostasis pathways. (a) Volcano plot of differentially expressed genes (DEGs) from RNA-seq of P1 hippocampus. Downregulated genes (blue) include ribosomal and translation-associated genes, while upregulated genes (red) are enriched for neurodevelopmental and synaptic function. FDR < 0.05, |log2FC| > 1. (b) Gene Ontology (GO) enrichment analysis of upregulated DEGs. Enriched biological processes include regulation of potassium ion transport, GABAergic synaptic transmission, dendritic spine morphogenesis, and neuron migration, indicating altered synaptic and neurodevelopmental programs. (c) Molecular function enrichment of upregulated DEGs. Overrepresented terms include high-voltage calcium channel activity, glutamate receptor activity, and GABA receptor binding, consistent with synaptic signaling alterations. (d) Cellular component enrichment of upregulated DEGs. Significant enrichment is observed in dendritic tree, synapse-associated complexes, glutamatergic synapse, and GABAergic synapse, suggesting structural remodeling of synaptic compartments. (e) GO enrichment of downregulated DEGs. Strongest enrichment is seen in translation and protein homeostasis categories, including cytoplasmic translation, ribosome biogenesis, chaperone-mediated protein folding, proteasome activity, mitochondrial ATP synthesis, and mRNA splicing. (f) STRING protein–protein interaction network of downregulated DEGs. Downregulated genes form interconnected modules centered on translation/ribosome complexes, spliceosome, proteasome, chaperonin-containing T-complex, mitochondrial respiratory chain complexes, and ATP-dependent chaperone networks. Additional modules include signaling clusters (Notch, Reelin, neuropeptide), highlighting widespread suppression of protein synthesis and proteostasis mechanisms in W300X-Het hippocampus.

Up-DEGs were highly represented in neurodevelopmental terms, such as regulations of neuroblast proliferation, neuron projection regulation, neuron generation, and cell migration (Fig. 5b-d, Supplemental Fig. 3a,b), correlated with the neurogenic and neuronal connectivity defects in W300X-Het hippocampus (Fig. 4). Further, critical genes involved in neuronal migration, axon guidance, and maturation, such as genes for Robo-Slit signaling, Fgf, Wnt, neurotrophin binding, and microtubule complex, were aberrantly upregulated (Fig. 5b-d).

The up-DEG set also revealed significant enrichment of synapse-related terms (Fig. 5b-d, Supplemental Fig. 3a,c), highlighting disruptions in synaptic organization, plasticity, and signaling pathways. Specifically, the following synapse-related terms were highly enriched in the upregulated gene set; dendritic spine morphogenesis and membrane depolarization, synapse assembly, chemical transmission, learning, memory, GABAergic and glutamatergic synapse signaling (e.g., *Gabrb1, Gabra3, Grm1*), and regulation of potassium, calcium, and chloride channels (e.g., *Kcnt1, Kcnt2, Kcnip2, Cacna1D, Scn5a, Cacna1e, Cacna1g*). Cadherin and integrin signaling genes (e.g., *Cdh22*, *Cdh24*, *Fat1*), linked to abnormal dendritic spine formation, were also upregulated in the W300X-Het hippocampus.

The down-DEG set also showed enrichment for neuronal and synaptic terms, including neuron fate specification, neuronal differentiation, spontaneous synaptic transmission, dendritic spine morphogenesis, and vesicle transport (Fig. 5e, Supplementary Fig. 3b,c, 4a,b). Notably, genes implicated in Reelin and Notch signaling pathways were downregulated (Fig. 5f), in line with their essential role in NSC maintenance, radial scaffold formation, neuronal positioning, and synaptic integration^32-34,36^.

These findings provide a comprehensive view of the early transcriptomic alterations in the W300X-Het hippocampus, highlighting that disrupted neurogenic and synaptic pathways underlie the cellular and structural defects observed at later stages.

### Protein homeostasis is disrupted W300X-Het hippocampus

Intriguingly, the W300X-Het hippocampus showed a significant downregulation across a spectrum of genes associated with core cellular functions, including protein translation, folding, and degradation (Fig. 5e,f, Supplemental Fig. 4). Notably, these disruptions are characterized by the simultaneous suppression of numerous genes within the same pathway or cellular process, rather than the loss of a single key component (Fig. 5f, Supplemental Fig. 4), underscoring the systemic nature of these molecular defects.

A predominant theme of down-DEGs across all databases is the dysregulation of translation. Strong enrichment in categories, such as cytoplasmic translation, ribosomal subunit, eukaryotic translation initiation factor 2 complex, and eukaryotic translation elongation factor 1 complex, highlights this pathway (Fig. 5f, Supplemental Fig. 4a-c). Significantly downregulated genes in this pathway include 39 ribosomal protein genes (e.g., *Rpl3*), eukaryotic translation elongation factor genes (e.g., *Eef1b2*), and eukaryotic translation initiation factor genes (e.g., *Eif4a1*). These results indicate marked impairment in ribosomal biogenesis and translation processes in the W300X-Het hippocampus.

The terms involving protein folding, such as post-translational protein folding, chaperone cofactor-dependent protein refolding, and chaperonin-containing T-complex, were also significantly enriched (Fig. 5e,f, Supplementary Fig. 4b). The downregulated genes in this pathway include genes for multiple subunits of the chaperonin-containing TCP1 complex (CCT or TRiC complex), *Cct2, Cct3, cct4, Cct6a, Cct7, Cct8*, and *Tcp1*. (Fig. 5f, Supplementary Fig. 4b,d). The TCP1 complex is a molecular chaperone complex essential for folding newly synthesized proteins, including cytoskeletal proteins such as actin and tubulin, thereby ensuring proper protein conformation, stability, and function^37^. Further, genes for heat shock proteins *Hspa4, Hspa8, Hspa9*, and *Hspe1,* which are molecular chaperones involved in protein folding, protein transport, and stress response^38^, were downregulated (Fig. 5f, Supplementary Fig. 4d). These results indicate the destabilized protein conformation and disrupted proteostasis in W300X-Het brains.

Other significantly enriched terms for the downregulated genes include proteasome degradation and parkin ubiquitin proteasomal system pathway (Fig. 5f, Supplementary Fig. 4d), underscoring the defective protein degradation process in the W300X-Het hippocampus. Genes in these categories include proteasome complex subunits, *Psma4, Psmab1, Psmab4, Psmc3*, *Psmc6*, and *Psmad4*, and a key component of the ubiquitin-proteasome system *Fbxw7*^39,40^.

Combined, the transcriptome analyses indicated that protein synthesis, the folding of newly synthesized peptides, and the degradation of misfolded proteins were impaired in the W300X-Het hippocampus. Importantly, our studies suggest the systemic and coordinated nature of FOXG1-directed gene regulation in establishing proteostasis in the developing hippocampus.

### The FOXG1-MYCN pathway promotes the expression of translation-involved genes

FOXG1 is primarily recognized as a transcriptional repressor^10^. Consistent with this view, the analysis revealed that established FOXG1 target genes, such as *Slit1* and *Slit*^10^, were upregulated in the W300X-Het hippocampus (Supplementary Fig. 3b). However, the regulatory mechanism through which FOXG1 is engaged in the transcriptional activation of the target genes remains less understood.

To define the transcriptional regulatory modes by which FOXG1 controls the expression of the down-DEG set, we performed the following analyses. Firstly, the downregulation of the genes in W300X-Het brains may occur exclusively secondarily due to the aberrant upregulation of FOXG1’s direct target genes. To ask if any down-DEGs directly recruit FOXG1 to their regulatory elements, we compared them with FOXG1 ChIP-seq datasets, which mapped FOXG1-binding genomic loci^10^. 41% of down-DEGs were associated with FOXG1-binding peaks, while 48% of up-DEGs were annotated to FOXG1 peaks (Fig. 6a). FOXG1 peaks associated with down-DEGs were located more in intergenic and promoter regions of the genes than FOXG1 peaks for up-DEGs (Fig. 6b). These results indicate that at least a subset of down-DEGs directly recruit FOXG1 in the genome.

**Figure 6.**
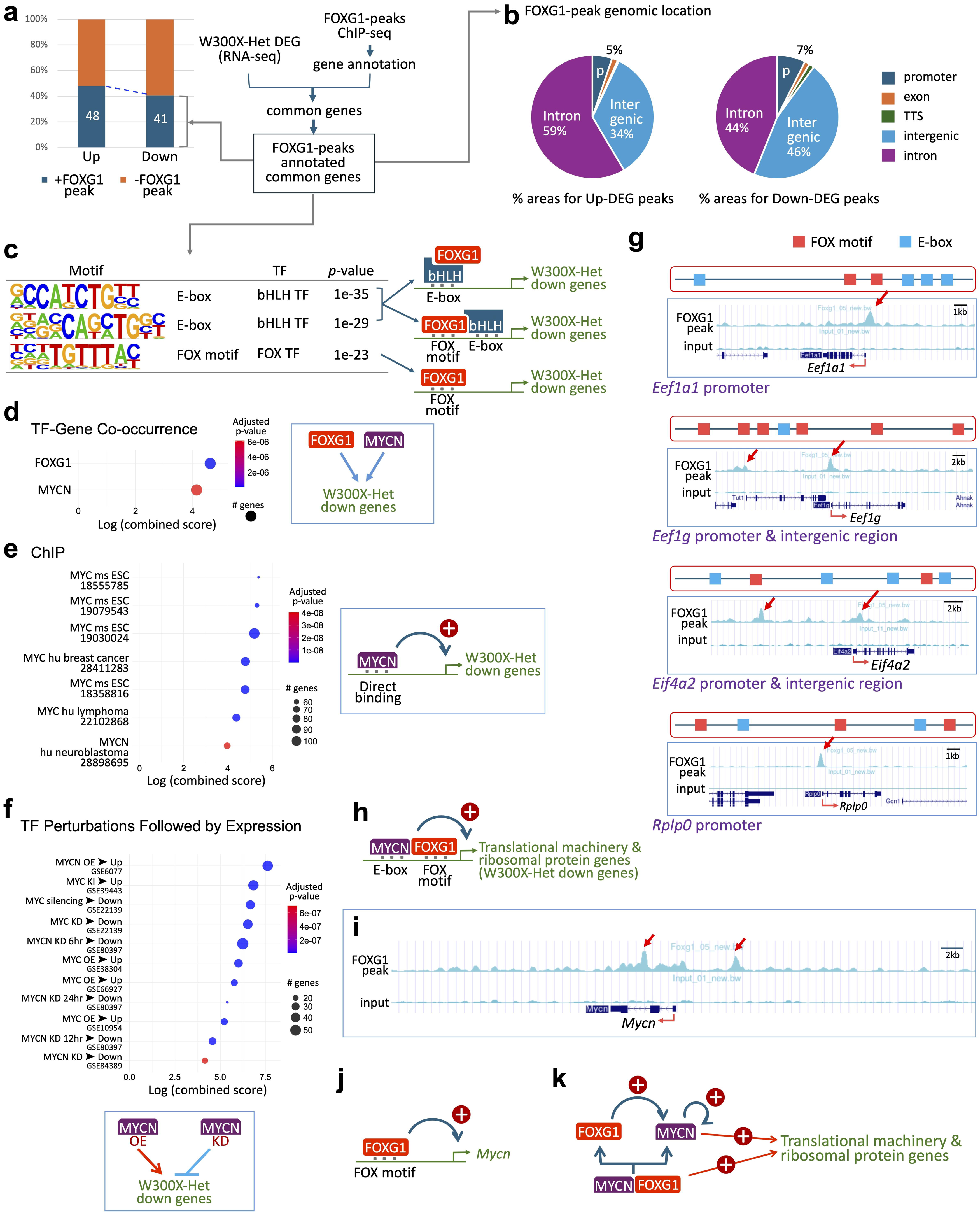
FOXG1-MYCN axis cooperatively regulates translation-associated genes in the W300X-Het hippocampus. (a) Integration of FOXG1 ChIP-seq with W300X-Het hippocampal RNA-seq identifies overlap between FOXG1-bound peaks and differentially expressed genes (DEGs). A substantial fraction of downregulated genes (41%) and upregulated genes (48%) are associated with FOXG1 peaks. (b) Genomic distribution of FOXG1-binding peaks reveals that down-DEG-associated peaks are enriched more in intergenic and promoter regions than up-DEG-associated peaks. (c) Motif enrichment analysis of FOXG1 peaks associated with down-DEGs shows strong enrichment of canonical FOX motifs and E-box motifs. These findings support a model in which FOXG1 and bHLH factors are co-recruited to a subset of FOXG1-binding genomic loci. (d) TF-gene co-occurrence analysis identifies MYCN as the most significantly associated bHLH TF linked to down-DEGs, suggesting a cooperative interaction between MYCN and FOXG1. (e) ChIP-X Enrichment Analysis (ChEA) of down-DEGs identifies MYC/MYCN as top transcription factors directly bound to down-DEG-associated peaks. (f) TF perturbation-expression signature analysis reveals that MYCN overexpression (OE) or knockdown (KD) strongly influences the W300X-Het down-DEG set, consistent with MYCN’s regulatory role. (g) FOXG1 ChIP-seq tracks at representative translation-associated genes (*Eef1a1*, *Eef1g*, *Eif4a2*, *Rplp0*) show co-occurrence of FOX and E-box motifs in FOXG1-bound peaks with both, indicating potential MYCN co-occupancy. (h) Schematic model of cooperative FOXG1-MYCN activation of translation machinery and ribosomal protein genes. FOXG1 and MYCN co-bind to overlapping loci, facilitating transcriptional activation. (i) FOXG1 ChIP-seq tracks reveal FOXG1-binding peaks in *Mycn* regulatory regions (promoter, intronic, intergenic), suggesting direct activation of *Mycn* transcription by FOXG1. (j) FOXG1 binding at the *Mycn* locus supports a feedforward mechanism in which FOXG1 sustains MYCN expression. (k) Proposed regulatory model. FOXG1 directly promotes *Mycn* expression and cooperatively binds with MYCN to translation-related gene loci. This FOXG1-MYCN transcriptional module amplifies ribosomal biogenesis and translational capacity.

Secondly, to explore how FOXG1 is recruited to these down-DEGs, we performed motif discovery analysis on FOXG1-binding peaks associated with these genes^41^. As expected, the FOX motif was enriched (Fig. 6c), indicating that FOXG1 binds to many down-DEGs by recognizing its own cognate element. Intriguingly, the most significantly enriched DNA signature was the two variations of the E-box motif, the binding site for the basic helix-loop-helix (bHLH) TF family (Fig. 6c), suggesting that bHLH TF may play a role in controlling a subset of down-DEGs by co-binding with FOXG1 or recruiting FOXG1 to their regulatory elements.

Thirdly, to identify TFs that are most likely involved in regulating the down-DEG set, we searched the “TF-Gene Co-occurrence” database using down-DEGs as input. This analysis revealed MYCN, the bHLH-leucine zipper TF, as the most significantly associated with down-DEGs among bHLH TFs (Fig. 6d).

Fourthly, to identify FOXG1’s partner TFs independently and unbiasedly, we employed the ChEA (ChIP-X Enrichment Analysis) database, which links TFs to their direct target genes based on experimental datasets of ChIP-seq or similar methods^42-44^. This analysis identified MYC and MYCN as the top candidates directly binding to many down-DEGs (Fig. 6e). Several ChIP-seq databases in multiple cell types, including human neuroblastoma^44-49^, showed that MYC or MYCN is recruited to the regulatory elements for down-DEGs in the W300X hippocampus. These results suggest that FOXG1 and MYC bind to the overlapping target genes.

Fifthly, we performed complementary analyses using the "TF Perturbations Followed by Expression" library, which links TFs with the signature gene expression pattern^50^. This approach allows us to pinpoint TFs whose altered expression leads to the coordinated upregulation or downregulation of genes within the down-DEG set, providing insight into the upstream regulators driving these molecular changes. Intriguingly, down-DEGs were highly associated with the gene expression changes caused by MYC overexpression or knockdown (Fig. 6f). The genes in the W300X-Het down-DEG set were either upregulated by the overexpression (OE) of MYCN or downregulated by the knockdown (KD) of MYCN, as determined by RNA-seq or similar transcriptome analyses in multiple cell types^51-58^ (Fig. 6f). These findings suggest a functional collaboration between FOXG1 and MYC/MYCN, where MYC and MYCN act as a key transcriptional partner in activating the genes belonging to the down-DEG set.

Among the three members of the MYC TF family, MYCN is particularly well-positioned to act as FOXG1’s partner in the hippocampus because it is highly expressed in the hippocampal DG and promotes the proliferation and maintenance of NPCs^59^. Furthermore, MYCN directly regulates genes involved in ribosomal biogenesis and the translation machinery^60,61^. These results suggest that FOXG1 and MYCN collaborate to control the translation-involved genes. To test if FOXG1 and MYCN bind to similar genomic areas for translation gene activation, we analyzed the sequences of FOXG1 ChIP-seq peaks annotated to translation-related genes. The FOXG1 peaks associated with translation genes contained both FOX and E-box motifs (Fig. 6g, Supplementary Fig. 5), suggesting that the bHLH TF binds to the same loci along with FOXG1. To test whether MYCN binds to the same genomic areas, we analyzed publicly available MYCN ChIP-seq datasets from human neuroblastoma cell^45^. MYCN was recruited to genomic regions overlapping with the FOXG1-binding loci associated with translation-related genes (Fig. 6g, Supplementary Fig. 6). These results suggest that FOXG1 and MYCN cooperatively activate translation-related genes by co-occupying their regulatory elements (Fig. 6h).

MYCN activates translation genes and sustains its own transcription partly via auto-regulatory feedback^62^. To test if FOXG1 directly regulates *Mycn* expression to enhance the FOXG1-MYCN axis, we determined FOXG1-binding to the *Mycn* gene in the FOXG1 ChIP-seq database. Interestingly, multiple FOXG1-binding peaks were identified in intronic and intergenic regions of the *Mycn* gene (Fig. 6i,j). Furthermore, *Mycn* expression was significantly downregulated in the W300X-Het hippocampus (Supplementary Fig. 4c), consistent with FOXG1’s role in maintaining MYCN levels.

Our findings reveal a dual role for FOXG1 in collaborating with MYCN to regulate translation genes (Fig. 6j,k). First, FOXG1 binds translation gene loci alongside MYCN, leveraging its partnership with MYCN to activate target gene expression. Second, FOXG1 directly promotes MYCN expression, establishing a regulatory axis that amplifies the transcription of translation-related genes. Together, our studies uncovered the FOXG1-MYCN pathway crucial for translation gene activation in the developing hippocampus. This model highlights FOXG1’s unexpected role as a transcriptional activator in the hippocampus, mediated through its interplay with MYCN.

### Cellular stress pathways are induced in the W300X-Het Hippocampus

The disruption of protein homeostasis, such as translational dysfunction, is a key factor in triggering cellular stress. Our transcriptomic analysis revealed a significant enrichment of cellular stress pathways in the W300X-Het hippocampus. In particular, stress-responsive pathways associated with glucocorticoid signaling were prominently activated, as evidenced by enrichment in databases for response to glucocorticoid and nuclear glucocorticoid receptor binding (Fig. 7a). These findings suggest glucocorticoid stress hormone pathway activation in the W300X-Het condition.

**Figure 7.**
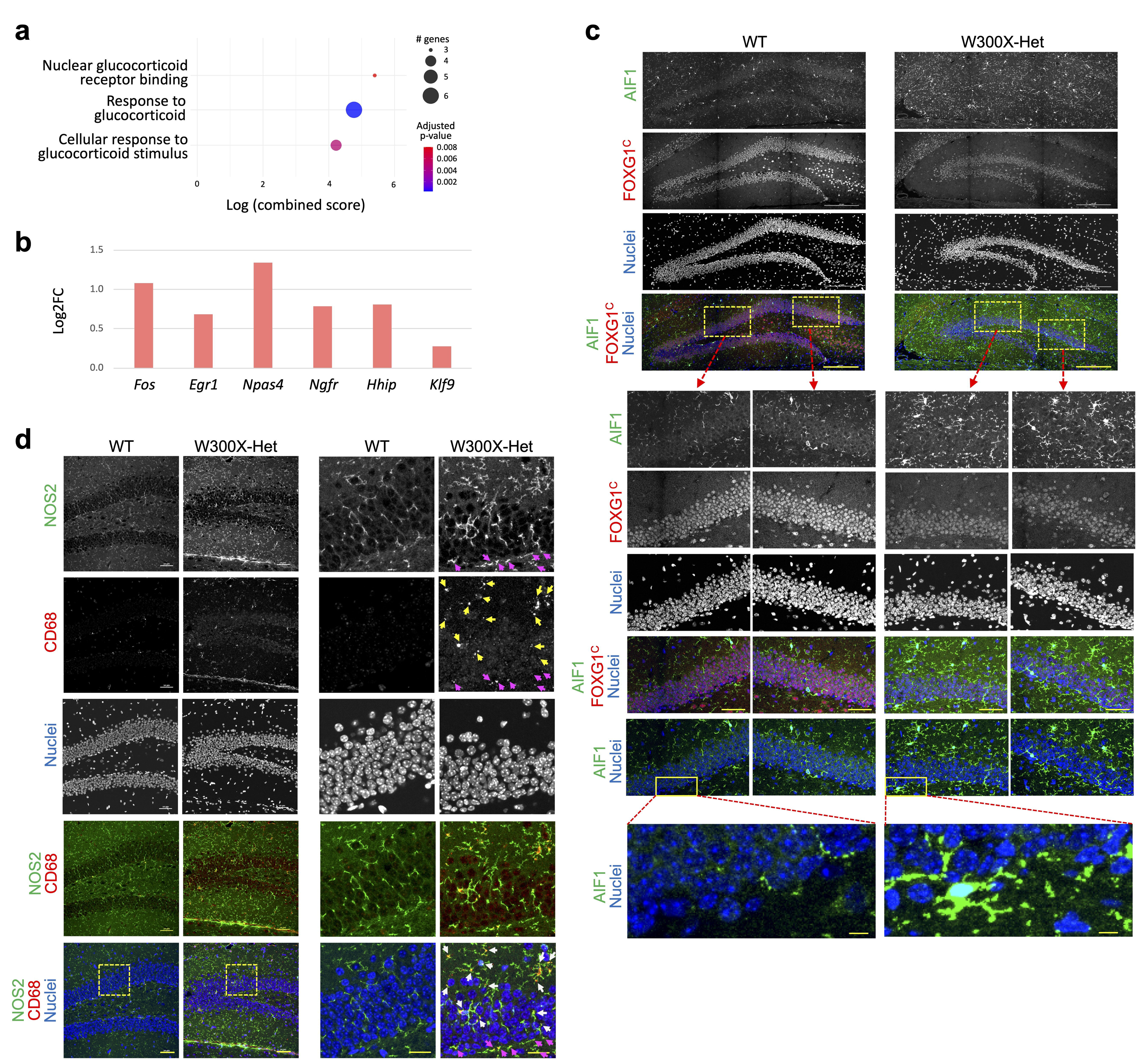
Cellular stress pathway activation and microglial response in W300X-Het dentate gyrus. (a) Pathway enrichment analysis of upregulated DEGs in P1 W300X-Het hippocampus reveals significant activation of nuclear glucocorticoid receptor binding and glucocorticoid-responsive pathways, consistent with induction of stress-response signaling. (b) Immediate early genes and stress-responsive transcripts are significantly upregulated in W300X-Het hippocampus, indicating transcriptional responses to cellular stress. (c) Immunostaining for AIF1 (IBA1) and FOXG1 in DG shows increased density and signal intensity of AIF1^+^ microglia in W300X-Het mice, with processes positioned adjacent to the subgranular zone (SGZ). Scale bars: 200 μm (top panels), 50 μm (middle panels), and 20 μm (bottom panels in a and b). (d) Co-immunostaining for NOS2 (iNOS) and CD68 shows pronounced increases in NOS2^+^ pro-inflammatory microglia and CD68^+^ lysosomal activity in W300X-Het DG. High-magnification images reveal colocalization of CD68 puncta (yellow arrows) within NOS2^+^ processes (white arrows), consistent with activated phagocytic microglia. The enrichment of activated microglia at the SGZ (magenta arrows) suggests potential non-cell-autonomous disruption of the neurogenic niche. Scale bars: 50 μm (left panels), and 20 μm (right panels with magnified images). (c,d) Each panel consists of paired columns (WT and W300X-Het) displaying images at the same magnification, with magnified regions outlined by yellow rectangles.

Further, several well-characterized stress-responsive genes were upregulated in the W300X-Het hippocampus (Fig. 7b). These included the immediate-early genes *Fos* and *Egr1*, which are rapidly induced by acute stress stimuli^63^. Other key stress-responsive genes, such as *Npas4, Klf9, Ngfr* (for p75NTR), and *Hhip* (for Hedgehog-interacting protein)^64-67^, were significantly upregulated. These genes respond to various forms of cellular stress, including neuronal excitotoxicity, inflammation, and glucocorticoid signaling, highlighting a broad activation of stress-response mechanisms in the W300X-Het hippocampus.

These findings demonstrate that cellular stress pathways are activated in the W300X-Het hippocampus. The upregulation of key stress-responsive genes suggests that the W300X mutation elicits a state of heightened vulnerability to cellular stress, potentially contributing to hippocampal dysfunction and impaired neurogenesis.

### Increased microglial activation in the DG of W300X-Het mice

Given the activation of cellular stress pathways in the W300X-Het hippocampus, we investigated whether neuroinflammation, particularly microglial activation, contributes to hippocampal dysfunction. To monitor microglial activation in the DG of W300X-Het mice, we performed IHC for microglial and inflammatory markers, AIF1 (aka IBA1), CD68, and NOS2 (aka iNOS for inducible nitric oxide synthase). AIF1, a calcium-binding protein expressed specifically in microglia^68^, was used to detect overall microglial presence and morphology. The density of AIF1^+^ cells was markedly elevated in W300X-Het mice compared to wild-type controls (Fig. 7c), indicating microglial proliferation or infiltration into the DG. Further, AIF1^+^ signal intensity was substantially stronger, and microglial processes were more readily visible in the W300X-Het DG (Fig. 7c), reflecting microglial activation. NOS2, a marker of pro-inflammatory microglia^69^, exhibited increased cell numbers and signal intensity in the W300X-Het DG (Fig. 7d). Similarly, CD68, a lysosomal marker associated with microglial phagocytosis^69^, showed a substantial increase in the W300X-Het DG (yellow arrows, Fig. 7d). Notably, fine, punctate speckled patterns of CD68 signals were located in NOS2^+^ processes (white arrows, Fig. 7d), indicating a transition of microglia into pro-inflammatory, phagocytic states in W300X-Het brains.

The activated microglia processes expressing AIF1, NOS2, and CD68 were aligned underneath the SGZ (Fig. 7c, magenta arrows, Fig. 7d), indicating that activated microglia are closely associated with the neurogenic niche of the DG. The spatial alignment between activated microglia and the SGZ suggests that inflammatory processes may modulate the local neurogenic microenvironment, thereby contributing to impaired neurogenesis in the W300X-Het hippocampus, given that microglia play crucial roles in maintaining stem cell niches under both homeostatic and pathological conditions^70^.

Overall, the findings suggest robust microglial activation and inflammation in the DG of W300X-Het mice.

### Learning and memory deficits in W300X-Het mice

Given the pronounced anatomical, cellular, and molecular abnormalities in the hippocampus of W300X-Het mice, we hypothesized that these defects would impair cognitive function. To test this idea, we subjected W300X-Het mice to the Morris Water Maze, a widely used test for assessing spatial learning and memory^71^ (Fig. 8a).

**Figure 8.**
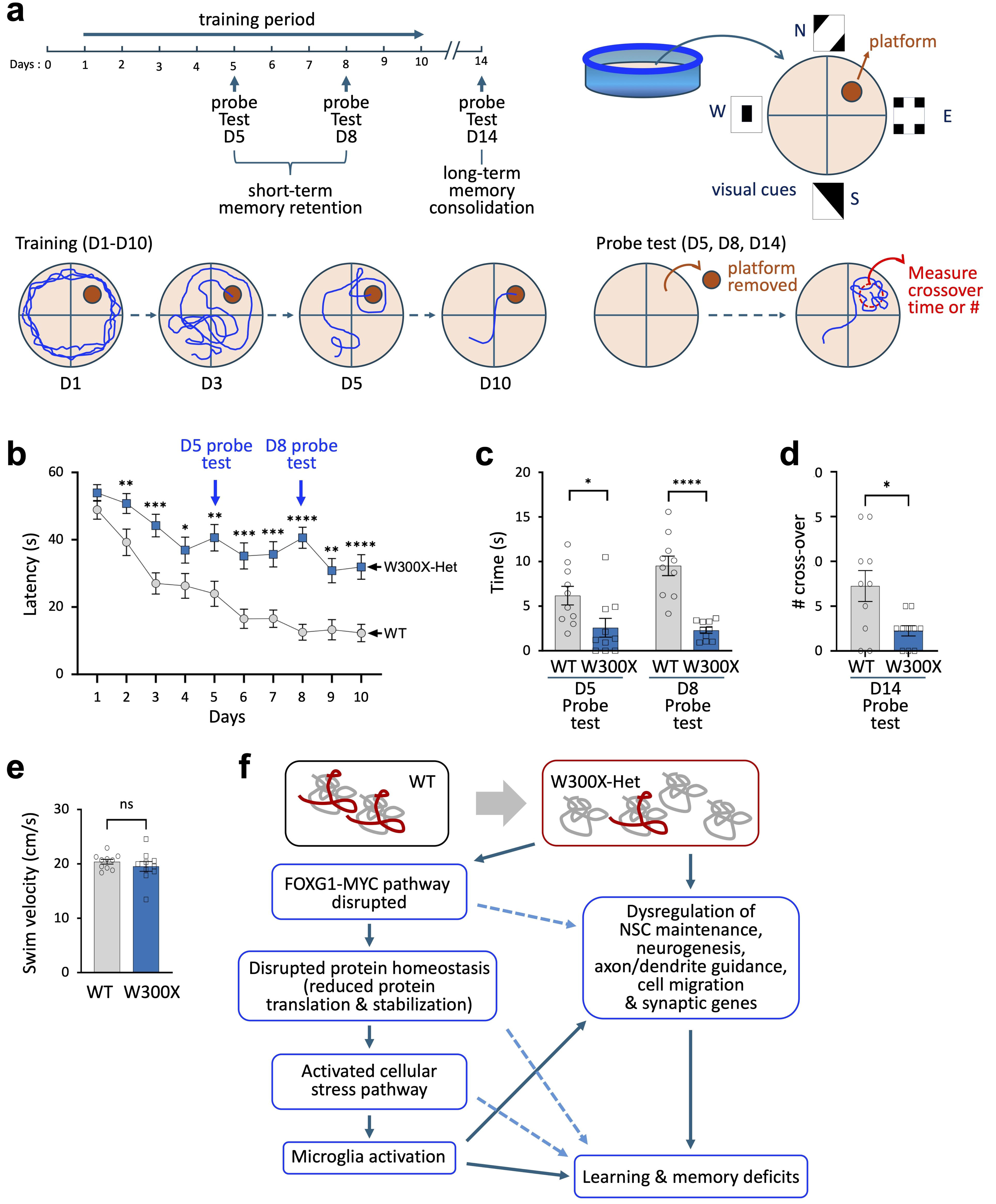
Spatial learning and memory impairments in W300X-Het mice and integrative pathogenic model. (a) Schematic of Morris water maze paradigm showing training (Days 1–10) and probe tests assessing short-term memory retention (Days 5 and 8) and long-term memory consolidation (Day 14). (b) W300X-Het mice display significantly prolonged escape latencies across the 10-day training period compared to WT controls, indicating impaired hippocampal-dependent learning. Note that training tests followed probe testing in D5 and D8. (c, d) In probe tests, W300X-Het mice show significantly reduced time spent in the target platform zone (D5, D8, c) and decreased platform crossover counts (D14, d), consistent with deficits in spatial memory encoding and retrieval. (e) Swim velocity is comparable between WT and W300X-Het mice, excluding motor impairments as a confound. (f) Integrative model. Truncated W300X protein exhibits enhanced stability and nuclear retention, disrupting FOXG1-MYCN-mediated activation of ribosomal biogenesis and translation machinery. Impaired protein homeostasis leads to reduced protein synthesis and destabilized proteostasis, which activates cellular stress pathways and microglial activation. Disruption of NSC maintenance, neurogenesis, axon/dendrite guidance, and synaptic gene programs, together with a pro-inflammatory DG environment, culminates in hippocampal dysfunction and severe learning and memory deficits in W300X-Het mice.

During the 10-day training period, W300X-Het mice exhibited significantly prolonged latencies in locating the submerged platform compared to WT controls throughout the training period (Fig. 8b). Notably, while WT mice displayed a steady and significant improvement in platform acquisition, W300X-Het mice showed delayed progression. The longer platform latency during training suggests impaired hippocampal-dependent learning processes.

In the first two probe tests that evaluate short-term memory retention at days 5 and 8, W300X-Het mice spent significantly less time within the platform proximity zone than WT mice (Fig. 8c). In the probe test performed 4 days after the training period, which evaluates long-term spatial memory consolidation at day 14, platform crossover counts were significantly reduced in W300X-Het mice (Fig. 8d). The reduced time spent in the target zone and fewer platform crossings in probe tests indicate that W300X-Het mice have difficulty encoding and retrieving spatial memory.

Notably, the swimming velocities of W300X-Het and WT mice were comparable (Fig. 8e), confirming that differences in cognitive performance were not confounded by motor impairments or motivational differences between the genotypes.

These findings provide robust evidence that W300X-Het mice suffer from substantial deficits in both spatial learning acquisition and spatial memory consolidation. Our study highlights the critical role of the FOXG1 gene in maintaining hippocampal function and spatial cognition.

## DISCUSSION

This study provides a comprehensive analysis of the W300X-Het mice, a patient-specific model of FS with a truncation-type FOXG1 variant, which unraveled the molecular and cellular mechanisms underlying the severe neurodevelopmental disorder FS (Fig. 8f). Our findings highlight profound disruptions in hippocampal development, protein homeostasis, and functional integrity, offering novel insights into the pathogenesis of FS. Furthermore, our results uncovered the FOXG1-MYCN pathway for hippocampal protein homeostasis and the interplay between FOXG1 mutations and neuroinflammatory pathways.

### Truncated FOXG1 Proteins in FS Pathogenesis

Our findings demonstrate that a truncation variant of the *FOXG1* gene, which belongs to the most prevalent type of FS-causing mutations, produces a stable fragment of the FOXG1 protein (Fig. 8f). The truncated W300X protein exhibits distinct features, including enhanced nuclear localization and increased stability compared to the full-length FOXG1 protein. These properties suggest that the W300X protein does not simply represent a loss-of-function mutation but exerts dominant-negative or gain-of-function effects. Future studies should explore the precise role of FOXG1 fragments in the pathogenic processes of FS, including their interactions with transcriptional coregulators and downstream targets. Such investigations would uncover therapeutic strategies targeting the activity of FOXG1 fragments, with implications for other truncation-type mutations in monogenic NDDs.

### Synaptic Dysfunction and Cognitive Impairments

The W300X-Het mice exhibit severe morphogenic defects in the hippocampus, including reduced NSC populations, deficient neurogenesis, and impaired granule cell migration and maturation. These findings align with prior studies on the loss-of-FOXG1 model, which shows reduced postnatal neurogenesis^23^. Interestingly, our study showed the W300X-Het DG phenotypes progressively worsen during postnatal stages, leading to marked structural reductions in adulthood, suggesting a compounded effect of NSC dysfunction, impaired neurogenic niches, and non-cell-autonomous mechanisms such as altered astrocytic support and microglial activation.

Structural abnormalities in the W300X-Het DG were accompanied by dysregulation of synapse-associated genes, suggesting significant synaptic dysfunction. Dysregulated expression of key genes involved in neurotransmitter signaling (e.g., GABAergic and glutamatergic synapses) and synaptic assembly (e.g., *Shank3, Grm1*) indicates impaired neural connectivity. These synaptic defects, combined with deficits in dendritic growth, branching, and neuron positioning, likely contribute to severely compromised neural circuits, where neurons fail to connect correctly, and synapses exhibit dysfunctional signaling. Behaviorally, W300X-Het mice exhibit pronounced spatial learning and memory impairments, which closely parallel hippocampal dysfunction in human FS patients. These findings underscore the utility of the W300X-Het model for understanding the structural and functional basis of cognitive deficits in FS.

### Protein Homeostasis and Ribosomal Stress

A striking feature of the W300X-Het hippocampus is the disruption of protein homeostasis, characterized by the downregulation of ribosomal and translational genes and impaired protein folding and degradation pathways. These deficits suggest a state of ribosomal stress, which compromises protein synthesis, neuronal growth, and synaptic formation^61,72^. The simultaneous reduction in chaperone and proteasome subunit expression exacerbates proteostatic dysfunction by impairing the folding and clearance of misfolded proteins, further contributing to cellular stress and neurodegeneration. Beyond transcriptional regulation, recent studies revealed that FOXG1 also modulates the microRNA processing pathway, mitochondrial dynamics, and the translation of select neuronal genes in cytosolic compartments^73-76^. Future studies are needed to explore how FOXG1’s nuclear transcriptional functions and cytosolic non-transcriptional roles are interconnected in maintaining protein homeostasis.

Interestingly, our findings implicate the FOXG1-MYCN pathway as a critical regulator of protein synthesis in the hippocampus, particularly in processes requiring robust translation, such as NSC division, synaptogenesis, and synaptic plasticity. FOXG1 directly promotes the expression of translation-associated genes by cooperating with MYCN, a master regulator of ribosomal biogenesis and protein synthesis^60,62^ (Fig. 6k). This collaboration is supported by the co-occurrence of FOXG1 and MYCN-binding motifs in translation gene promoters and the enrichment of MYCN target genes among FOXG1-bound transcriptional targets. The dual role of FOXG1 in regulating MYCN expression and collaborating with MYCN in activating target genes underscores a transcriptional network essential for these protein synthesis-dependent processes in the maturing hippocampus.

The FOXG1-MYCN pathway also raises intriguing parallels with cancer biology, as MYCN is a well-established oncogene driving high-risk neuroblastoma and medulloblastoma^77-79^. FOXG1 overexpression in medulloblastoma, where it promotes progenitor cell proliferation^80^, suggests potential roles for the FOXG1-MYCN axis in tumor biology. Understanding the interplay between FOXG1 and MYCN in both hippocampal development and tumorigenesis may uncover shared mechanisms linking neurodevelopmental disorders and cancer.

### NDDs and Neuroinflammation

Emerging evidence highlights the central role of neuroinflammation in NDDs, including ASD and Rett syndrome^81-83^. Microglial activation is a key mediator of neuroinflammation, releasing pro-inflammatory cytokines such as TNF-α and IL-1β, which can exacerbate neuronal dysfunction and cell death^84^. In the W300X-Het model, we observed robust microglial activation in the DG, as indicated by increased IBA1, CD68, and iNOS expression. These markers indicate a pro-inflammatory, phagocytic microglial phenotype, which impairs NSC maintenance and adult neurogenesis, thereby exacerbating the neurogenic and structural deficits.

The spatial association of activated microglia with the SGZ raises the possibility that inflammatory signals disrupt the neurogenic niche. Persistent microglial overactivation, driven by misfolded proteins or translational dysfunction, may create a chronic inflammatory environment, exacerbating neurodevelopmental deficits. Therapeutic strategies targeting microglial activation or cytokine signaling could, therefore, mitigate inflammation-driven hippocampal dysfunction in FOXG1-related disorders.

### Integrative Pathogenic Mechanisms

This study establishes a mechanistic framework linking FOXG1 mutations to molecular, cellular, and behavioral deficits characteristic of FS. By integrating transcriptomic and functional analyses, we identified protein homeostasis and neuroinflammation as central mediators of disease pathology. These findings suggest several therapeutic targets, including strategies to restore proteostasis via translational machinery or chaperone enhancement and to modulate neuroinflammation through microglial inhibitors or anti-cytokine therapies. By advancing our understanding of the molecular mechanisms underlying FS, these efforts could inform the development of targeted interventions to improve outcomes for FS individuals.

## METHOD

### Animals

All animal experimental procedures were conducted in accordance with the guidelines implemented by the Institutional Animal Care and Use Committee at University at Buffalo. W300X mutant and wild-type littermate mice were maintained on a background of C57BL6/J strain. All animals were housed in University at Buffalo animal facility. Animals were allowed ad libitum access to food and water and were reared on a 12h light-12h dark cycle.

W300X mutant mice were generated in C57BL/6J background by Crispr/Cas9-mediated gene targeting. The mutation c.900G>A was introduced by single guide RNA, Cas9 protein, and single stranded donor oligonucleotides for homology directed repair process. Guide RNA sequence was 5’-GGA GCC GGC GCG GTC CAT GA-3’, and donor oligonucleotides was 5’-GCC GCT CCA CCA CGT CTC GGG CCA AGC TGG CCT TTA AGC GCG GGG CGC GCC TCA CCT CCA CCG GCC TCA CAT TCA TGG ACC GCG CCG GAT CCC TCT ACT GAC CCA TGT CGC CCT TCC TGT CCC TGC ACC ACC CCC GCG CCA GCA GCA CTT T-3’ (plus strand), which includes restriction enzyme BamHI recognition site for the genotyping of offspring. Potential founder mice were screened by Sanger sequencing and restriction enzyme digestion for PCR product with primers flanking the target site. The targeted founder mice were backcrossed to C57BL/6J mice for at least five generations to eliminate potential off-target mutations before any experiments.

### Brain preparation, immunohistochemical analysis, and image acquisition

Dissected brains were fixed in 4% paraformaldehyde in phosphate-buffered saline (PBS) at 4°C overnight, equilibrated in 30% sucrose, and embedded in Tissue Freezing Medium^TM^ (Electron Microscopy Sciences) for frozen sectioning for 8 μm brain sections using cryostat (CM1950, Leica). Position matched sections were then stained following the standard immunohistochemical method. Sections were incubated with the primary antibody and the secondary antibodies conjugated to fluorophores (Jackson ImmunoResearch), and then counterstained with DAPI to reveal nuclei. Images were collected using Eclipse Ti2 confocal microscope (analyzed by Nikon Software). The primary antibodies included guinea pig anti-FOXG1^C^ (homemade), rabbit anti-FOXG1^N^ (Invitrogen #PA5-41493, 1:1000), mouse anti-GFAP (Millipore Sigma MAB360, 1:1000), rabbit anti-SOX2 (Millipore Ab5602, 1:1000), rabbit anti-TBR2 (Abcam ab183991, 1:1000), rabbit anti-MKI67 (Abcam ab15580, 1:1000), rabbit anti-MCM2 (Abcam ab108935, 1:1000), rabbit anti-PROX1(ANGIOBIO 11-002p, 1:1000), rabbit anti-NEUN (Abcam 177487, 1:1000), rabbit anti-DCX (Cell Signaling Technology #4604, 1:1000), goat anti-AIF1(Abcam, ab5076, 1:1000), mouse anti-NOS2 (Invitrogen MA5-17139, 1:1000), and rat anti-CD68(Abcam ab53444, 1:1000) antibodies. The secondary antibodies from Jackson ImmunoResearch Laboratories included Alexa Fluor 488 donkey anti-guinea pig (706-545-148), Cy3 donkey anti-rabbit (711-165-152), Cy3 donkey anti-rat (712-165-150), Cy3 donkey anti-guinea pig(706-165-148), Alexa Fluor 488 donkey anti-mouse(715-545-150), and Alexa Fluor 488 donkey anti-rabbit (711-545-152) antibodies.

High resolution images (60X magnification, tiles) were employed in CellProfiler for the automated quantification of immunofluorescent intensity in individual hippocampal cells. To detect the FOXG1 antibody immunostaining signal in individual cells, the IdentifyPrimaryObjects module (employing Global thresholding) and the Otsu approach (based on object intensity settings) were employed. The settings were adjusted to capture all positive signals while minimizing the noise. The MeasureObjectIntensity module was used to generate intensity data for further analysis. The Export to Spreadsheet module was applied to export all data into Excel format for subsequent processing in Prism.

### Western blot analysis

Mouse cortex and hippocampus were homogenized in RIPA buffer (Cell Signaling Technology, #9806) supplemented with Protease Inhibitor Cocktail (Cell Signaling Technology, #7012). An aliquot of the cortical and hippocampal homogenates was used to determine total protein concentrations using BCA kit (ThermoPierce). Fifteen micrograms of protein were loaded onto 10–20% Bis-Tris Novex gels (Life Technologies) for electrophoresis. Proteins were then transferred to PVDF membranes using the iBlot system (Life Technologies). Western blotting was performed with 5% BSA blocking solution and primary antibodies, rabbit anti-FOXG1^C^ (Abcam, ab196868, 1:1000) and rabbit anti-FOXG1^N^ (Invitrogen, #PA5-41493, 1:1000). Secondary antibodies were applied, and membranes were imaged and quantified using Odyssey LI-COR equipment and software (LI-COR Biosciences).

### RNA-seq data analyses

Total RNA from hippocampal tissues at the P1 stage were isolated from both hemispheres and extracted by PureLink RNA Mini Kit (Invitrogen, 12183025) according to the manufacture’s protocol. RNA-seq analyses were performed using hippocampus of W300X-Het and WT mice (n=3 for W300X-Het mice; n=3 for WT). We checked the quality of the raw reads using FastQC. Clean reads were mapped to the mouse reference genome (mm10) using STAR (v2.6.1d)^85^ with default parameters. RSEM (v1.3.3)^86^ was used to quantify the gene expression levels and the DEGs were identified using EBseq^87^. The absolute value of log2 fold change of 0.3 and false discovery rate (FDR) of 0.05 were set as criteria to determine if the change of expression level of a gene was significant. Gene set enrichment analysis (GSEA) was subsequently carried out using Enrichr^43^ on the upregulated DEGs and the downregulated DEGs, separately.

### ChIP-seq Data Analyses

For mouse FOXG1 ChIP-seq data^10^, FastQC was used to check the quality of the raw reads. Clean reads were aligned to the mouse reference genome (mm10) using BWA-mem (0.7.17-r1188)^88^ with default parameters. The mapped reads were filtered and sorted using samtools (v1.8)^89^ and the duplicated reads were marked by MarkDuplicates of Picard tools (v2.27.4) (http://broadinstitute.github.io/picard). The reads were indexed by samtools for the further analyses. Callpeak of MACS2 (v2.2.7.1)^90^ was used to call the significantly enriched peaks compared to the control samples. The peaks were annotated and motif analysis on the peaks was carried out using annotatePeaks and findMotifsGenome of HOMER (v4.11.1) ^41^, respectively. We downloaded human MYCN ChIP-seq data (GSE83728 and GSE94822) from Gene Expression Omnibus (GEO) and analyzed them to compare with the mouse FOXG1 ChIP-seq data to discover the partnership of FOXG1 and MYCN. The reads were mapped to human reference genome (hg38) and analyzed by the same pipeline we used for the mouse data. Due to the species difference, we used BLAT (v39x1)^91^ to search the similar regions in the human genome to the FOXG1-binding regions in the mouse genome based on the mouse reference genome sequence where the FOXG1 peaks were located. Then we compared the regions to the MYCN-binding regions to see if there was any association between FOXG1 and MYCN.

### Morris Water Maze Test

The arena of Morris water maze has diameter of 120 cm filled with preheated 21℃ water. Non-toxic white paint was dissolved in the water prior to the experiment to make water opaque so the platform is not visible to the mice. The Morris water maze arena was surrounded by black screens that had distinctive white visual marks on four different sides. Movement of the mice was recorded and measured by Noldus Ethovision XT. The Morris water maze test was performed during the light phase and all mice were habituated for at least 60 minutes before the test each day. The Morris water maze test was performed consecutively from day 1 to day 10 of learning periods while the transparent platform was shallowly submerged for 1 cm at a fixed location. Four trials of the learning test were performed each day, with 10 minutes intervals between each trial. A mouse was placed on one of four quadrants at randomly predetermined orders and allowed to swim for 60 seconds or until the mouse found the platform. Then, the mouse was placed on the platform for 20 seconds to allow the mouse to learn spatial information of the surroundings after each trial. Probe tests were performed on day 5, 8 and 14 before the learning test of the day without the platform in the arena.

Probe tests were run for 60 seconds for a single trial a day.

## Acknowledgments

We are grateful to Hyeryeong Park, Taeyeon Kim, Muhua Liu, and Xuefang He for their various contributions to this study. Also, we greatly appreciate the efforts and support of the staff and veterinarians of the University at Buffalo LAF. This work was funded by grants from NINDS/NIH (NS111760 and NS100471 to S.-K.L. and NS118748 to S.-K.L. and J.W.L), FOXG1 Research Foundation (to S.-K.L. and J.W.L) and Simons Foundation (SFARI 1013720 to S.-K.L.) as well as the generous startup fund from University at Buffalo (to S.-K.L. and J.W.L).

## Author Contributions

S.J., J.W.L. and S.-K.L. designed the experiments and interpreted the data. J.M. designed and performed the computational data analysis. S.J., L.L., D.S., J.P., and E. K. carried out the experiments and analyzed the data. J.W.L. and S.-K. L. supervised the work.

