## Supplemental figures for "Human patient-specific FOXG1 syndrome mouse model revealed FOXG1-MYCN-mediated regulation of protein homeostasis in neurodevelopmental disorder"

### Slide 1
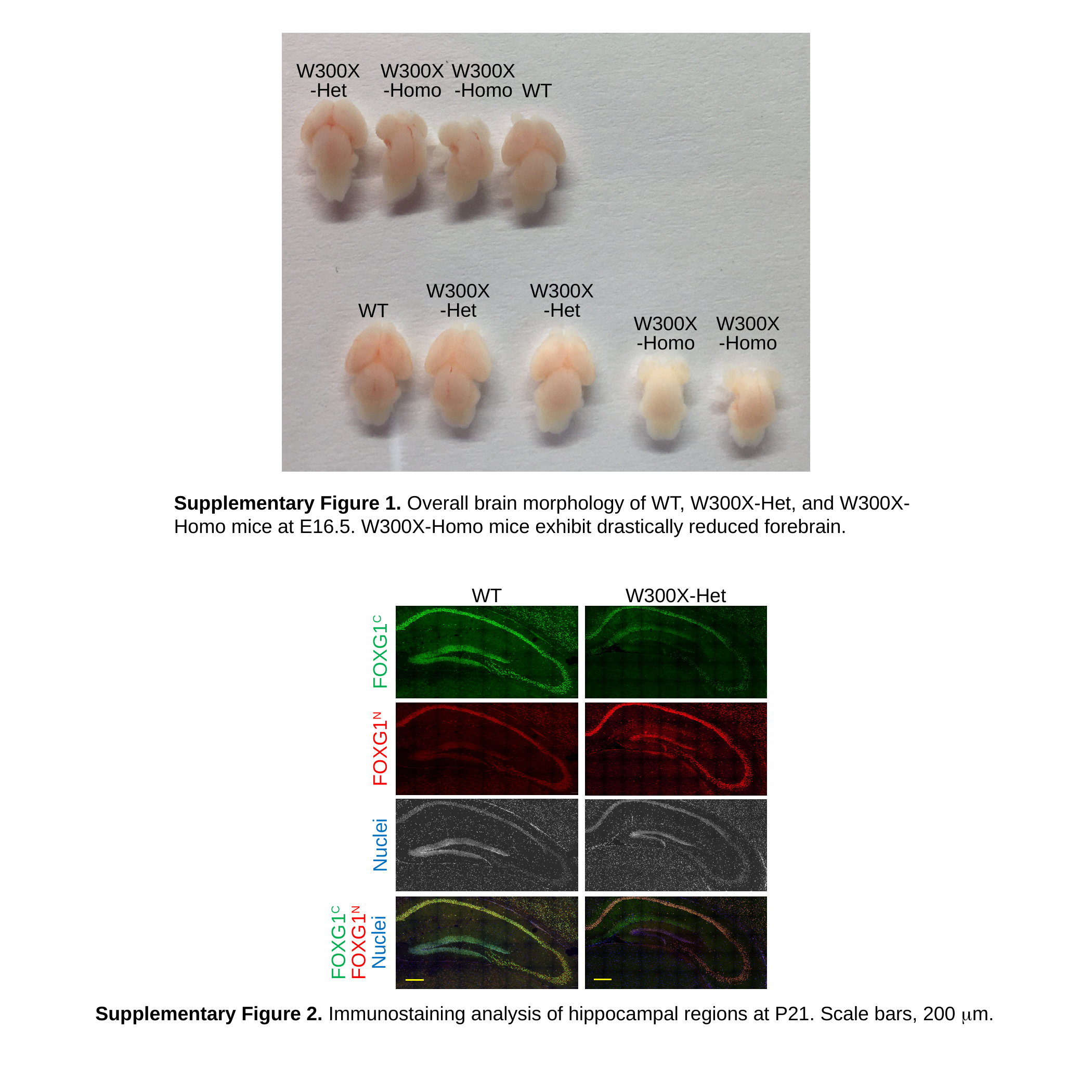

W300X
-Het
W300X
-Homo
W300X
-Homo
WT
W300X
-Het
W300X
-Het
WT
W300X
-Homo
W300X
-Homo
Supplementary Figure 1. Overall brain morphology of WT, W300X-Het, and W300X-Homo mice at E16.5. W300X-Homo mice exhibit drastically reduced forebrain.
WT
W300X-Het
FOXG1^C^
FOXG1^N^
Nuclei
FOXG1^C^
FOXG1^N^
Nuclei
Supplementary Figure 2. Immunostaining analysis of hippocampal regions at P21. Scale bars, 200 μm.

### Slide 2
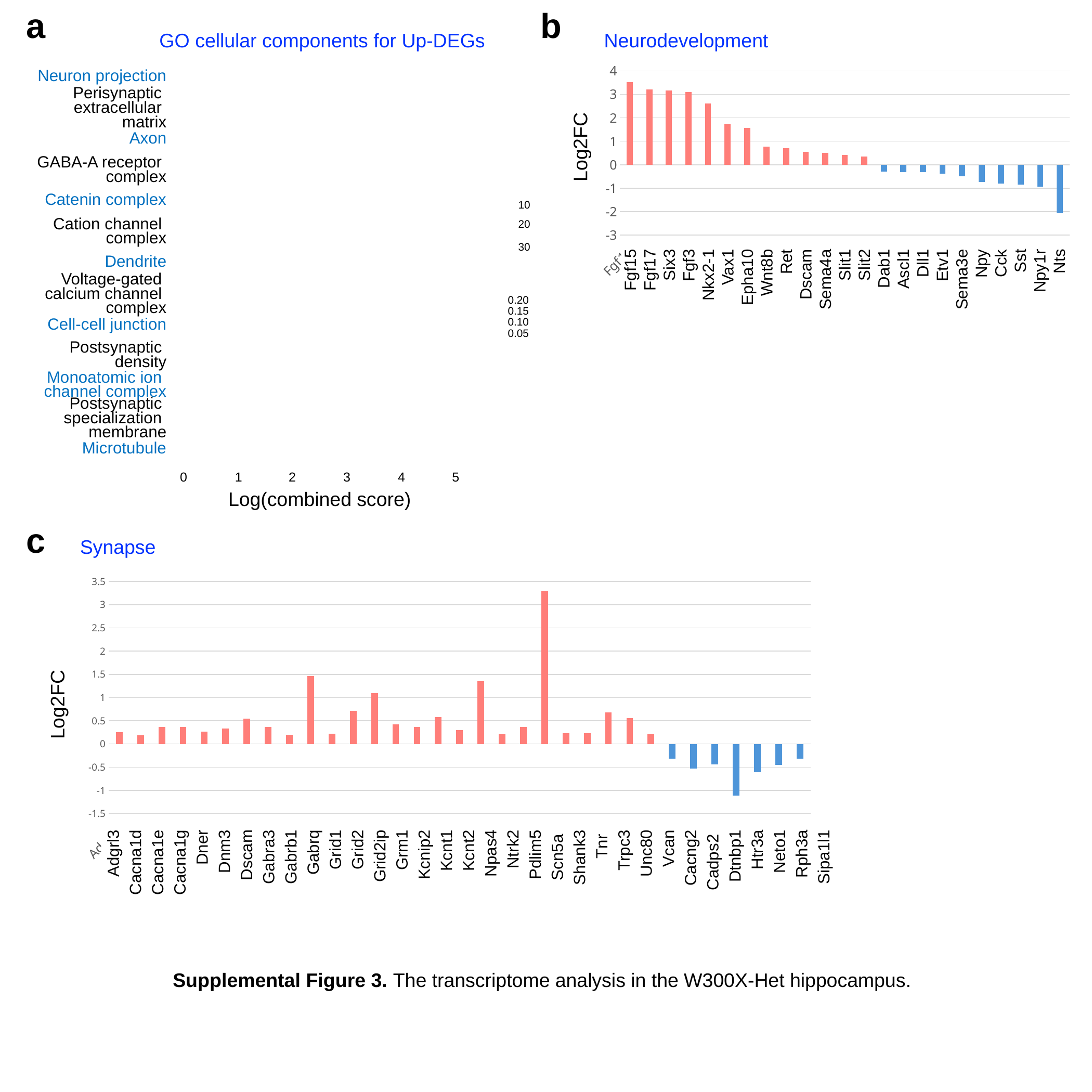

a
b
GO cellular components for Up-DEGs
Neurodevelopment
#### Chart
| Category | |
|---|---|
| Fgf15 | 3.5184884005572044 |
| Fgf17 | 3.2049851961254903 |
| Six3 | 3.1611086055850532 |
| Fgf3 | 3.1018296733002177 |
| Nkx2-1 | 2.605476021599403 |
| Vax1 | 1.7579006466945286 |
| Epha10 | 1.5698841749865633 |
| Wnt8b | 0.7714562268138896 |
| Ret | 0.7103888195127378 |
| Dscam | 0.5433232143951773 |
| Sema4a | 0.5021033524174457 |
| Slit1 | 0.4247649 |
| Slit2 | 0.355524324 |
| Dab1 | -0.2837388712339288 |
| Ascl1 | -0.31485045741641826 |
| Dll1 | -0.3179147315606107 |
| Etv1 | -0.39226123653202555 |
| Sema3e | -0.4964710332703343 |
| Npy | -0.7381170326040568 |
| Cck | -0.8005427931820263 |
| Sst | -0.8409630945454604 |
| Npy1r | -0.9266374262784971 |
| Nts | -2.0715095894022753 |Fgf15
Fgf17
Six3
Fgf3
Nkx2-1
Vax1
Epha10
Wnt8b
Ret
Dscam
Sema4a
Slit1
Slit2
Dab1
Ascl1
Dll1
Etv1
Sema3e
Npy
Cck
Sst
Npy1r
Nts
Neuron projection
Perisynaptic
extracellular
matrix
Log2FC
Axon
GABA-A receptor
complex
Catenin complex
10
20
Cation channel
complex
30
Dendrite
Voltage-gated
calcium channel
complex
0.20
0.15
0.10
Cell-cell junction
0.05
Postsynaptic
density
Monoatomic ion
channel complex
Postsynaptic
specialization
membrane
Microtubule
0
1
2
3
4
5
Log(combined score)
Adgrl3
Cacna1d
Cacna1e
Cacna1g
Dner
Dnm3
Dscam
Gabra3
Gabrb1
Gabrq
Grid1
Grid2
Grid2ip
Grm1
Kcnip2
Kcnt1
Kcnt2
Npas4
Ntrk2
Pdlim5
Scn5a
Shank3
Tnr
Trpc3
Unc80
Vcan
Cacng2
Cadps2
Dtnbp1
Htr3a
Neto1
Rph3a
Sipa1l1
c
Synapse
#### Chart
| Category | |
|---|---|
| Adgrl3 | 0.25723692795208974 |
| Cacna1d | 0.18812744400499873 |
| Cacna1e | 0.3663310589858118 |
| Cacna1g | 0.3686300463197297 |
| Dner | 0.26550326763640786 |
| Dnm3 | 0.3359165096989565 |
| Dscam | 0.5433232143951773 |
| Gabra3 | 0.3644403083754569 |
| Gabrb1 | 0.20192940051756364 |
| Gabrq | 1.459215718636324 |
| Grid1 | 0.21572022895021747 |
| Grid2 | 0.7081424644997566 |
| Grid2ip | 1.0987587746331002 |
| Grm1 | 0.4254725127368306 |
| Kcnip2 | 0.36807809811261843 |
| Kcnt1 | 0.5825158497334293 |
| Kcnt2 | 0.29616593205312985 |
| Npas4 | 1.347922074296566 |
| Ntrk2 | 0.20474527737864553 |
| Pdlim5 | 0.36740419439461247 |
| Scn5a | 3.2880389176750744 |
| Shank3 | 0.22840566017314387 |
| Tnr | 0.23370440872223286 |
| Trpc3 | 0.6826284524097409 |
| Unc80 | 0.5516723626249466 |
| Vcan | 0.2046443583817731 |
| Cacng2 | -0.31221360583796903 |
| Cadps2 | -0.5307570224031506 |
| Dtnbp1 | -0.44429582347492136 |
| Htr3a | -1.1101757453540921 |
| Neto1 | -0.6050668599758927 |
| Rph3a | -0.4519572801884065 |
| Sipa1l1 | -0.3154060398052951 |Log2FC
Supplemental Figure 3. The transcriptome analysis in the W300X-Het hippocampus.

### Slide 3
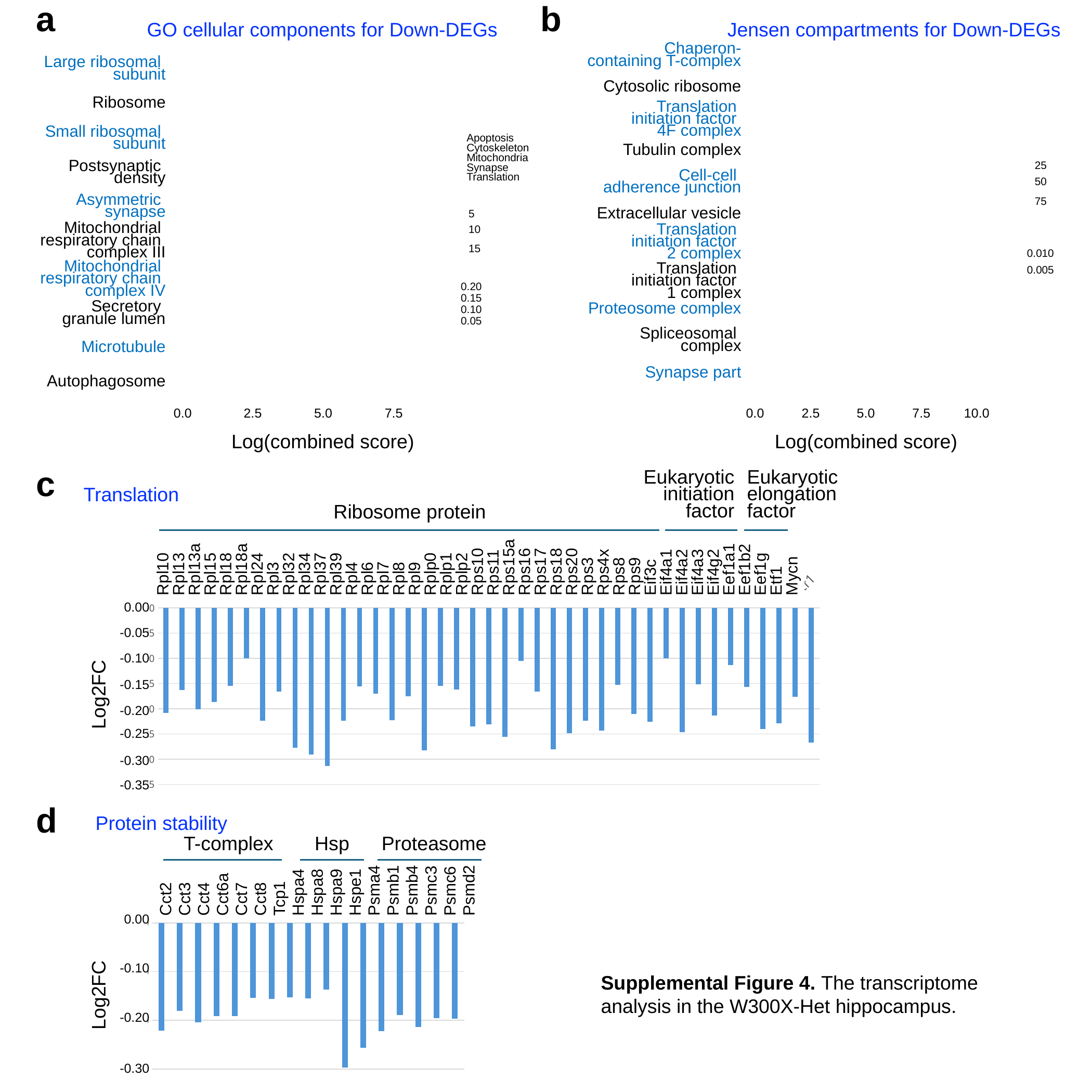

a
b
GO cellular components for Down-DEGs
Jensen compartments for Down-DEGs
Chaperon-
containing T-complex
Large ribosomal
subunit
Cytosolic ribosome
Ribosome
Translation
initiation factor
4F complex
Small ribosomal
subunit
Apoptosis
Cytoskeleton
Mitochondria
Synapse
Translation
Tubulin complex
25
Postsynaptic
density
Cell-cell
adherence junction
50
75
Asymmetric
synapse
5
Extracellular vesicle
10
Mitochondrial
respiratory chain
complex III
Translation
initiation factor
2 complex
Rpl10
Rpl13
Rpl13a
Rpl15
Rpl18
Rpl18a
Rpl24
Rpl3
Rpl32
Rpl34
Rpl37
Rpl39
Rpl4
Rpl6
Rpl7
Rpl8
Rpl9
Rplp0
Rplp1
Rplp2
Rps10
Rps11
Rps15a
Rps16
Rps17
Rps18
Rps20
Rps3
Rps4x
Rps8
Rps9
Eif3c
Eif4a1
Eif4a2
Eif4a3
Eif4g2
Eef1a1
Eef1b2
Eef1g
Etf1
Mycn
15
0.010
Mitochondrial
respiratory chain
complex IV
0.005
Translation
initiation factor
1 complex
0.20
0.15
0.10
Secretory
granule lumen
Proteosome complex
0.05
Spliceosomal
complex
Microtubule
Synapse part
Autophagosome
0.0
2.5
5.0
7.5
0.0
2.5
5.0
7.5
10.0
Log(combined score)
Log(combined score)
c
Eukaryotic initiation factor
Eukaryotic elongation factor
Translation
Ribosome protein
#### Chart
| Category | log2FC |
|---|---|
| Rpl10 | -0.20817675902297508 |
| Rpl13 | -0.16252270083791787 |
| Rpl13a | -0.20081741655078467 |
| Rpl15 | -0.18694902267274197 |
| Rpl18 | -0.15511558081972082 |
| Rpl18a | -0.10005895165505138 |
| Rpl24 | -0.22326424874550418 |
| Rpl3 | -0.16644191327160013 |
| Rpl32 | -0.27743484999676515 |
| Rpl34 | -0.29034920452551694 |
| Rpl37 | -0.31388912209644765 |
| Rpl39 | -0.2236931635066739 |
| Rpl4 | -0.15529882591387176 |
| Rpl6 | -0.16962847445709914 |
| Rpl7 | -0.22259907047704025 |
| Rpl8 | -0.1756096795345422 |
| Rpl9 | -0.28247513607827995 |
| Rplp0 | -0.15480130137436876 |
| Rplp1 | -0.1621127799797876 |
| Rplp2 | -0.2347980609346792 |
| Rps10 | -0.23086713832042127 |
| Rps11 | -0.25521831395029204 |
| Rps15a | -0.104796397481855 |
| Rps16 | -0.16625195303229143 |
| Rps17 | -0.2799601307201695 |
| Rps18 | -0.24832886570112775 |
| Rps20 | -0.22355161057268957 |
| Rps3 | -0.24277297663199932 |
| Rps4x | -0.1523752199419284 |
| Rps8 | -0.20999825372141748 |
| Rps9 | -0.22596561117490324 |
| Eif3c | -0.10027816980855273 |
| Eif4a1 | -0.24654373650295947 |
| Eif4a2 | -0.15112507707502423 |
| Eif4a3 | -0.21342151337154303 |
| Eif4g2 | -0.11290131231620706 |
| Eef1a1 | -0.15662376560928193 |
| Eef1b2 | -0.24047423102517085 |
| Eef1g | -0.2283258766138616 |
| Etf1 | -0.1763312910728649 |
| Mycn | -0.2671816878007206 |0.00
-0.05
-0.10
-0.15
Log2FC
-0.20
Cct2
Cct3
Cct4
Cct6a
Cct7
Cct8
Tcp1
Hspa4
Hspa8
Hspa9
Hspe1
Psma4
Psmb1
Psmb4
Psmc3
Psmc6
Psmd2
-0.25
-0.30
-0.35
d
Protein stability
T-complex
Hsp
Proteasome
#### Chart
| Category | |
|---|---|
| Cct2 | -0.22147414533266874 |
| Cct3 | -0.18104460668025005 |
| Cct4 | -0.20362487977345106 |
| Cct6a | -0.19175897604897815 |
| Cct7 | -0.191835418541363 |
| Cct8 | -0.15440441911204214 |
| Tcp1 | -0.15627560466047657 |
| Hspa4 | -0.15261002020781653 |
| Hspa8 | -0.15543044219314409 |
| Hspa9 | -0.1371477807557674 |
| Hspe1 | -0.29725084202027424 |
| Psma4 | -0.25601642589172036 |
| Psmb1 | -0.22275272459359133 |
| Psmb4 | -0.18951800795034293 |
| Psmc3 | -0.21324783906518843 |
| Psmc6 | -0.19551653204391373 |
| Psmd2 | -0.196636722277648 |
0.00
-0.10
Supplemental Figure 4. The transcriptome analysis in the W300X-Het hippocampus.
Log2FC
-0.20
-0.30

### Slide 4
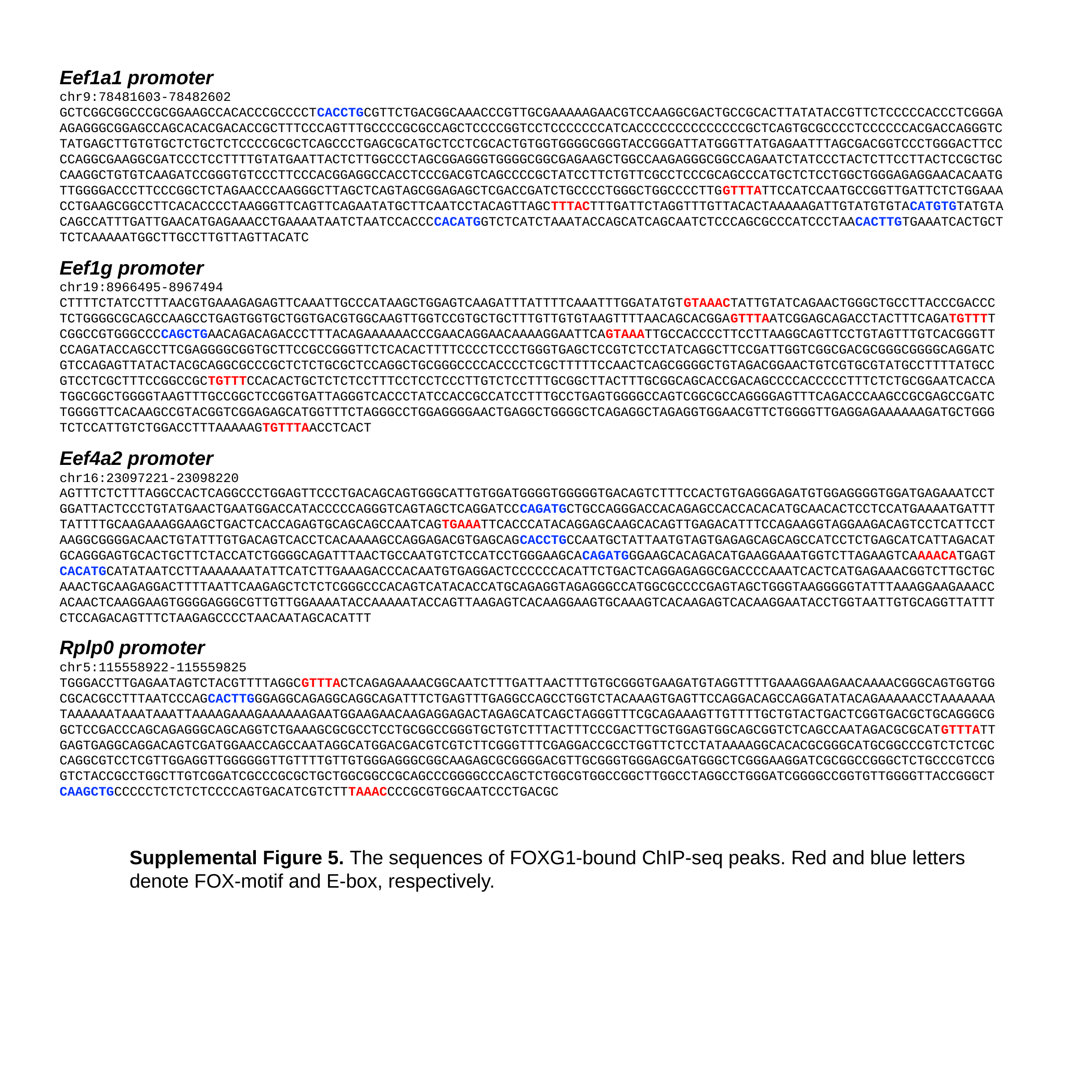

Eef1a1 promoter
chr9:78481603-78482602
GCTCGGCGGCCCGCGGAAGCCACACCCGCCCCTCACCTGCGTTCTGACGGCAAACCCGTTGCGAAAAAGAACGTCCAAGGCGACTGCCGCACTTATATACCGTTCTCCCCCACCCTCGGGAAGAGGGCGGAGCCAGCACACGACACCGCTTTCCCAGTTTGCCCCGCGCCAGCTCCCCGGTCCTCCCCCCCATCACCCCCCCCCCCCCCGCTCAGTGCGCCCCTCCCCCCACGACCAGGGTCTATGAGCTTGTGTGCTCTGCTCTCCCCGCGCTCAGCCCTGAGCGCATGCTCCTCGCACTGTGGTGGGGCGGGTACCGGGATTATGGGTTATGAGAATTTAGCGACGGTCCCTGGGACTTCCCCAGGCGAAGGCGATCCCTCCTTTTGTATGAATTACTCTTGGCCCTAGCGGAGGGTGGGGCGGCGAGAAGCTGGCCAAGAGGGCGGCCAGAATCTATCCCTACTCTTCCTTACTCCGCTGCCAAGGCTGTGTCAAGATCCGGGTGTCCCTTCCCACGGAGGCCACCTCCCGACGTCAGCCCCGCTATCCTTCTGTTCGCCTCCCGCAGCCCATGCTCTCCTGGCTGGGAGAGGAACACAATGTTGGGGACCCTTCCCGGCTCTAGAACCCAAGGGCTTAGCTCAGTAGCGGAGAGCTCGACCGATCTGCCCCTGGGCTGGCCCCTTGGTTTATTCCATCCAATGCCGGTTGATTCTCTGGAAACCTGAAGCGGCCTTCACACCCCTAAGGGTTCAGTTCAGAATATGCTTCAATCCTACAGTTAGCTTTACTTTGATTCTAGGTTTGTTACACTAAAAAGATTGTATGTGTACATGTGTATGTACAGCCATTTGATTGAACATGAGAAACCTGAAAATAATCTAATCCACCCCACATGGTCTCATCTAAATACCAGCATCAGCAATCTCCCAGCGCCCATCCCTAACACTTGTGAAATCACTGCTTCTCAAAAATGGCTTGCCTTGTTAGTTACATC
Eef1g promoter
chr19:8966495-8967494
CTTTTCTATCCTTTAACGTGAAAGAGAGTTCAAATTGCCCATAAGCTGGAGTCAAGATTTATTTTCAAATTTGGATATGTGTAAACTATTGTATCAGAACTGGGCTGCCTTACCCGACCCTCTGGGGCGCAGCCAAGCCTGAGTGGTGCTGGTGACGTGGCAAGTTGGTCCGTGCTGCTTTGTTGTGTAAGTTTTAACAGCACGGAGTTTAATCGGAGCAGACCTACTTTCAGATGTTTTCGGCCGTGGGCCCCAGCTGAACAGACAGACCCTTTACAGAAAAAACCCGAACAGGAACAAAAGGAATTCAGTAAATTGCCACCCCTTCCTTAAGGCAGTTCCTGTAGTTTGTCACGGGTTCCAGATACCAGCCTTCGAGGGGCGGTGCTTCCGCCGGGTTCTCACACTTTTCCCCTCCCTGGGTGAGCTCCGTCTCCTATCAGGCTTCCGATTGGTCGGCGACGCGGGCGGGGCAGGATCGTCCAGAGTTATACTACGCAGGCGCCCGCTCTCTGCGCTCCAGGCTGCGGGCCCCACCCCTCGCTTTTTCCAACTCAGCGGGGCTGTAGACGGAACTGTCGTGCGTATGCCTTTTATGCCGTCCTCGCTTTCCGGCCGCTGTTTCCACACTGCTCTCTCCTTTCCTCCTCCCTTGTCTCCTTTGCGGCTTACTTTGCGGCAGCACCGACAGCCCCACCCCCTTTCTCTGCGGAATCACCATGGCGGCTGGGGTAAGTTTGCCGGCTCCGGTGATTAGGGTCACCCTATCCACCGCCATCCTTTGCCTGAGTGGGGCCAGTCGGCGCCAGGGGAGTTTCAGACCCAAGCCGCGAGCCGATCTGGGGTTCACAAGCCGTACGGTCGGAGAGCATGGTTTCTAGGGCCTGGAGGGGAACTGAGGCTGGGGCTCAGAGGCTAGAGGTGGAACGTTCTGGGGTTGAGGAGAAAAAAGATGCTGGGTCTCCATTGTCTGGACCTTTAAAAAGTGTTTAACCTCACT
Eef4a2 promoter
chr16:23097221-23098220 AGTTTCTCTTTAGGCCACTCAGGCCCTGGAGTTCCCTGACAGCAGTGGGCATTGTGGATGGGGTGGGGGTGACAGTCTTTCCACTGTGAGGGAGATGTGGAGGGGTGGATGAGAAATCCTGGATTACTCCCTGTATGAACTGAATGGACCATACCCCCAGGGTCAGTAGCTCAGGATCCCAGATGCTGCCAGGGACCACAGAGCCACCACACATGCAACACTCCTCCATGAAAATGATTTTATTTTGCAAGAAAGGAAGCTGACTCACCAGAGTGCAGCAGCCAATCAGTGAAATTCACCCATACAGGAGCAAGCACAGTTGAGACATTTCCAGAAGGTAGGAAGACAGTCCTCATTCCTAAGGCGGGGACAACTGTATTTGTGACAGTCACCTCACAAAAGCCAGGAGACGTGAGCAGCACCTGCCAATGCTATTAATGTAGTGAGAGCAGCAGCCATCCTCTGAGCATCATTAGACATGCAGGGAGTGCACTGCTTCTACCATCTGGGGCAGATTTAACTGCCAATGTCTCCATCCTGGGAAGCACAGATGGGAAGCACAGACATGAAGGAAATGGTCTTAGAAGTCAAAACATGAGTCACATGCATATAATCCTTAAAAAAATATTCATCTTGAAAGACCCACAATGTGAGGACTCCCCCCACATTCTGACTCAGGAGAGGCGACCCCAAATCACTCATGAGAAACGGTCTTGCTGCAAACTGCAAGAGGACTTTTAATTCAAGAGCTCTCTCGGGCCCACAGTCATACACCATGCAGAGGTAGAGGGCCATGGCGCCCCGAGTAGCTGGGTAAGGGGGTATTTAAAGGAAGAAACCACAACTCAAGGAAGTGGGGAGGGCGTTGTTGGAAAATACCAAAAATACCAGTTAAGAGTCACAAGGAAGTGCAAAGTCACAAGAGTCACAAGGAATACCTGGTAATTGTGCAGGTTATTTCTCCAGACAGTTTCTAAGAGCCCCTAACAATAGCACATTT
Rplp0 promoter
chr5:115558922-115559825 TGGGACCTTGAGAATAGTCTACGTTTTAGGCGTTTACTCAGAGAAAACGGCAATCTTTGATTAACTTTGTGCGGGTGAAGATGTAGGTTTTGAAAGGAAGAACAAAACGGGCAGTGGTGGCGCACGCCTTTAATCCCAGCACTTGGGAGGCAGAGGCAGGCAGATTTCTGAGTTTGAGGCCAGCCTGGTCTACAAAGTGAGTTCCAGGACAGCCAGGATATACAGAAAAACCTAAAAAAATAAAAAATAAATAAATTAAAAGAAAGAAAAAAGAATGGAAGAACAAGAGGAGACTAGAGCATCAGCTAGGGTTTCGCAGAAAGTTGTTTTGCTGTACTGACTCGGTGACGCTGCAGGGCGGCTCCGACCCAGCAGAGGGCAGCAGGTCTGAAAGCGCGCCTCCTGCGGCCGGGTGCTGTCTTTACTTTCCCGACTTGCTGGAGTGGCAGCGGTCTCAGCCAATAGACGCGCATGTTTATTGAGTGAGGCAGGACAGTCGATGGAACCAGCCAATAGGCATGGACGACGTCGTCTTCGGGTTTCGAGGACCGCCTGGTTCTCCTATAAAAGGCACACGCGGGCATGCGGCCCGTCTCTCGCCAGGCGTCCTCGTTGGAGGTTGGGGGGTTGTTTTGTTGTGGGAGGGCGGCAAGAGCGCGGGGACGTTGCGGGTGGGAGCGATGGGCTCGGGAAGGATCGCGGCCGGGCTCTGCCCGTCCGGTCTACCGCCTGGCTTGTCGGATCGCCCGCGCTGCTGGCGGCCGCAGCCCGGGGCCCAGCTCTGGCGTGGCCGGCTTGGCCTAGGCCTGGGATCGGGGCCGGTGTTGGGGTTACCGGGCTCAAGCTGCCCCCTCTCTCTCCCCAGTGACATCGTCTTTAAACCCCGCGTGGCAATCCCTGACGC
Supplemental Figure 5. The sequences of FOXG1-bound ChIP-seq peaks. Red and blue letters denote FOX-motif and E-box, respectively.

### Slide 5
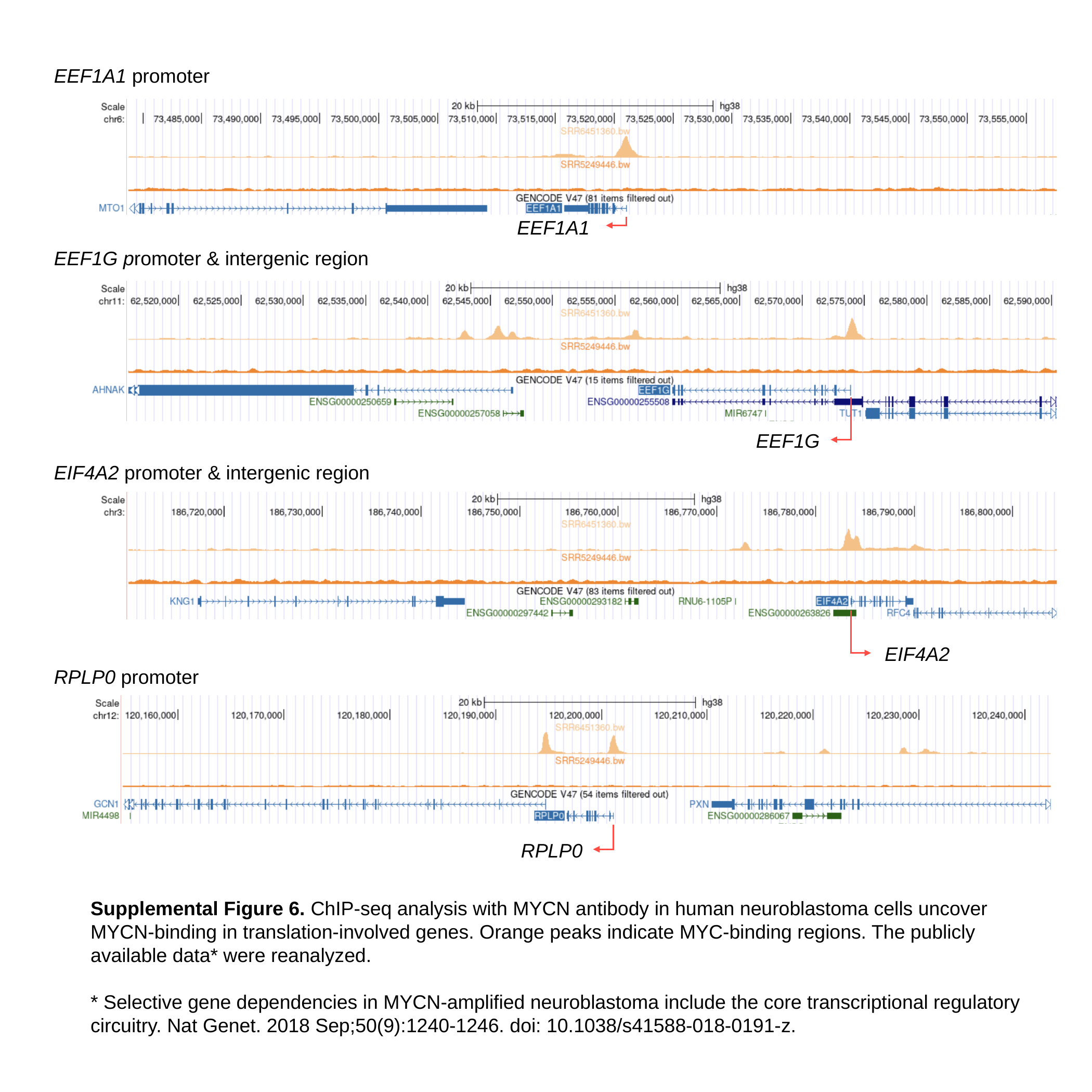

EEF1A1 promoter
EEF1A1
EEF1G promoter & intergenic region
EEF1G
EIF4A2 promoter & intergenic region
EIF4A2
RPLP0 promoter
RPLP0
Supplemental Figure 6. ChIP-seq analysis with MYCN antibody in human neuroblastoma cells uncover MYCN-binding in translation-involved genes. Orange peaks indicate MYC-binding regions. The publicly available data* were reanalyzed.
* Selective gene dependencies in MYCN-amplified neuroblastoma include the core transcriptional regulatory circuitry. Nat Genet. 2018 Sep;50(9):1240-1246. doi: 10.1038/s41588-018-0191-z.
